# Discovery of imidazole-based inhibitors of *P. falciparum* cGMP-dependent protein kinase

**DOI:** 10.1101/2021.11.05.467463

**Authors:** Rammohan R. Yadav Bheemanaboina, Mariana Laureano de Souza, Mariana Lozano Gonzalez, Shams Ul Mahmood, Tyler Eck, Tamara Kreiss, Samantha O. Aylor, Alison Roth, Patricia Lee, Brandon S. Pybus, Dennis J. Colussi, Wayne E. Childers, John Gordon, John J. Siekierka, Purnima Bhanot, David P. Rotella

## Abstract

The discovery of new targets for treatment of malaria and in particular those aimed at the pre-erythrocytic stage in the life cycle, advanced with the demonstration that orally administered inhibitors of *Plasmodium falciparum* cGMP-dependent protein kinase (PfPKG) could clear infection in a murine model. This enthusiasm was tempered by unsatisfactory safety and/or pharmacokinetic issues found with these chemotypes. To address the urgent need for new scaffolds, this manuscript presents initial structure-activity relationships in an imidazole scaffold at four positions, representative *in vitro* ADME, hERG characterization and cell-based anti-parasitic activity. This series of PfPKG inhibitors has good *in vitro* PfPKG potency, low hERG activity and cell-based anti-parasitic activity against multiple *Plasmodium* species that appears to correlate with *in vitro* potency.

---

The emergence of artemisinin-resistant *Plasmodium falciparum* in Africa^1,2^ and the slowing decline in deaths from malaria^3^ signal the need to identify new targets for prophylaxis and treatment.^**4**^ The pre-erythrocytic portion of the life cycle is an attractive and comparatively under explored point for therapeutic attack because of the very low parasite burden compared to other life cycle stages. Drugs that target these stages are an essential component of the anti-malarial effort because a decrease in liver infection by sporozoites significantly reduces severity and incidence of malaria.^6^ There is a comparative paucity of candidates in this area of anti-malarial drug development.^5,7^

To address this issue, *Plasmodium falciparum* cGMP-dependent protein kinase (PfPKG) is of particular interest because it is essential in pre-erythrocytic, asexual and sexual stages of the parasite.^8,9,10^ Baker and co-workers described the discovery and optimization of an orally bioavailable, potent, selective small molecule inhibitor (**1**, Figure 1). This imidazopyridine cleared infection at a dose of 10 mg/kg orally in a SCID mouse model.^11,12,13^

**Figure 1:**
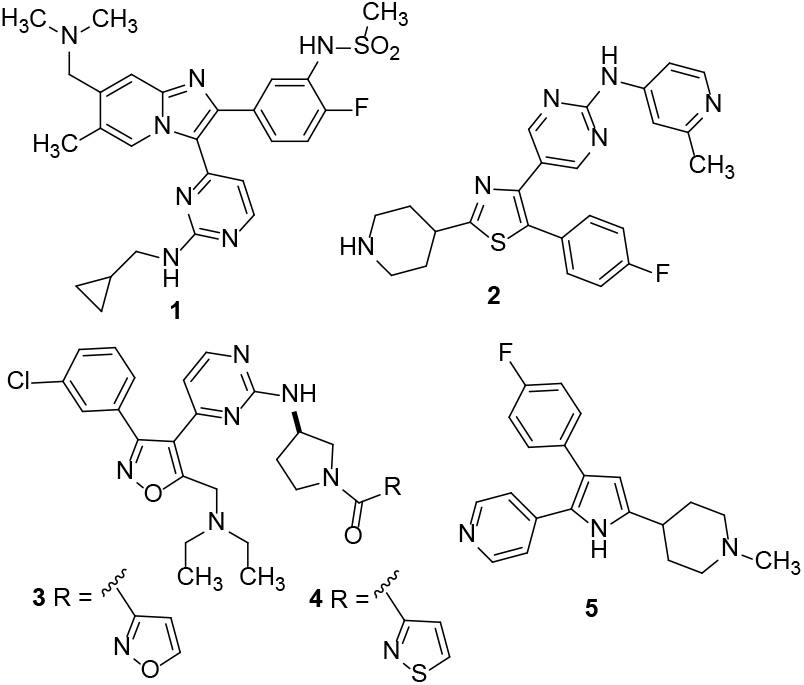
PfPKG Inhibitors

A subsequent report that unspecified examples in this series were Ames positive and limited progression of the scaffold.^14^ The Baker group disclosed trisubstituted thiazoles such as **2** (Figure 1) that exhibited rapid killing of *P. falciparum* in culture.^15^ Interestingly this desirable property was independent of PfPKG inhibition. Proteomic experiments suggested that inhibition of a serine/arginine protein kinase SRPK2 was a key contributor to rapid parasite killing, comparable to artesunate, a recognized standard. Examples in this series of thiazoles, including **2**, showed single digit micromolar hERG activity and/or *in vitro* metabolic instability limiting their use in more advanced studies.

Important pharmaceutical property and safety issues can be addressed by identifying new chemical matter to provide novel candidates to address this unmet need. We previously reported the discovery of an isoxazole chemotype, exemplified by **3** and **4** (Figure 1).^16^ These compounds showed enzymatic potency comparable (IC_50_s ∼20 nM) to known pyr-role **5**^17^ *in vitro* against PfPKG, and were not active against human PKG or the T618Q mutant PfPKG^18^ at 10 μM. Parasite expressing this mutant enzyme demonstrate lower sensitivity to PfPKG inhibitors such as **5** but retain enzymatic activity and the ability to proceed through the life cycle ^18^.

In the course of the focused screen that identified isoxazole hits, we also identified an imidazole scaffold **6** (Figure 2) that was active in a PfPKG screening assay. We were attracted to the comparatively low molecular weight of this chemotype and the potential for optimization at multiple positions that could result in improved properties. Additionally, the synthesis of this class of compounds was significantly shorter than the isoxazole series.

**Figure 2:**
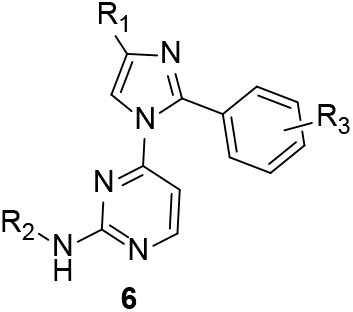
Imidazole PfPKG

Based on previous experience, we chose to focus first on exploration of the amino substituent (R_2_) on the pyrimidine. The synthesis of this set of derivatives is outlined in Scheme 1. Commercially available 4-methyl-2-phenyl imidazole was deprotonated with sodium hydride then treated with 4-chloro-2-methythiopyrimidine in dry DMF at 60-70°C to arylate the imidazole nitrogen, followed by oxone-mediated conversion to the sulfone. Displacement of the sulfone with a variety of diamines, Boc-deprotection and acylation with preferred carboxylic acids as described previously^16^ afforded the target amides **10a**-**h**.

It is evident from the data in Table 1 that the (R)-3-aminopyrrolidine linker is strongly preferred compared to the other cyclic amine variations, a cyclopropyl alkyl diamine and the S-enantiomer of 3-aminopyrrolidine. We previously observed this stereoselective effect in the isoxazole class of PfPKG inhibitors.^16^ In this case, all alternative amine linkers afforded inactive PfPKG inhibitors whereas with the isoxazole scaf-fold-selected examples retained some activity.^16^ This suggests distinct and specific SAR at this point in the structure. An additional difference in SAR for the imidazole scaffold is the more pronounced potency difference between the 2-thiazolyl and 1-N-methyl-3-pyrazolyl amides, with a strong preference for the former. In this group, the most potent derivative (**10a**, PfPKG IC_50_ 320 nM) underwent further evaluation to provide baseline data for this chemotype on selectivity for human PKG and the T618Q mutant PfPKG. We observed that **10a** had excellent selectivity versus these related PKGs (3-10% inhibition @ 10 μM).

**Scheme 1:**
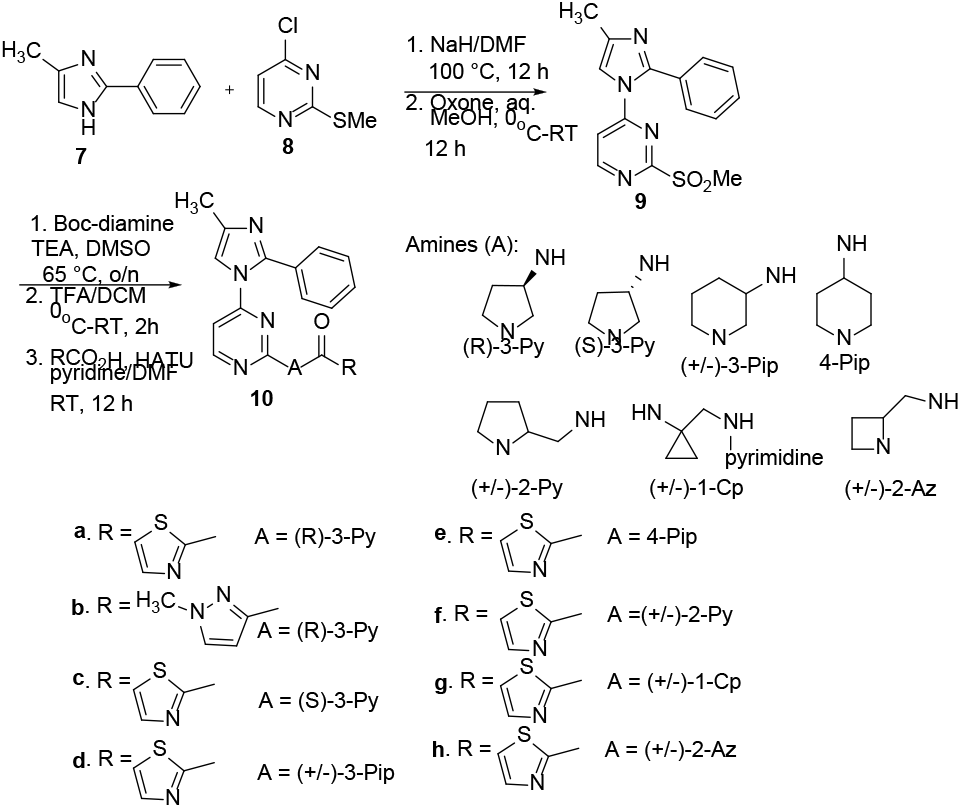
4-methyl imidazole PfPKG Inhibitors

**Table 1:**
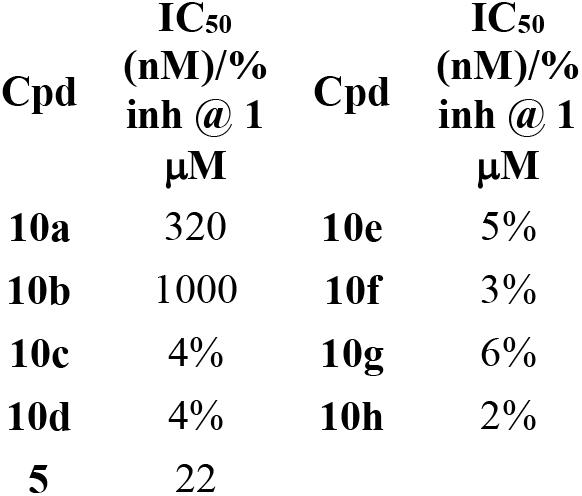
*in vitro* PfPKG inhibition by 4-methyl imidazoles **10a**-**10h**

| Cpd | IC<sub>50</sub><br>(nM)/%<br>inh @ 1<br>μM | Cpd | IC<sub>50</sub><br>(nM)/%<br>inh @ 1<br>μM |
| --- | --- | --- | --- |
| **10a** | 320 | **10e** | 5% |
| **10b** | 1000 | **10f** | 3% |
| **10c** | 4% | **10g** | 6% |
| **10d** | 4% | **10h** | 2% |
| **5** | 22 |  |  |

Assessment of *in vitro* ADME characteristics revealed that **10a** was metabolically unstable in human (HLM) and murine (MLM) liver microsomes (half life less than 2 minutes) with excellent water solubility (200 μM) and moderate CYP3A4 inhibition (IC_50_ 0.94 μM). We elected to address the metabolic stability issue first by replacing the 4-methyl group and substituting the aromatic ring because we viewed these as potential site(s) for oxidative metabolism. The 4-methyl group was replaced with a cyclopropyl ring and in view of existing isoxazole SAR with phenyl substituents^16^, chose to target an un-substituted and 3-chlorophenyl derivative. The synthesis, outlined in Scheme 2, condensed appropriate chlorobenzamidines with cyclopropyl bromomethyl ketone **12** to afford 4-cyclopropyl-2-phenyl imidazoles **13a** and **13b** in good yield. Following the steps outlined in Scheme 1, the targets **14a** and **14b** were obtained in a straightforward manner. In parallel, using the 4-methylimidazole template, we explored a sampling of aryl substituents on the benzene ring to explore this feature of SAR. The synthesis of these analogs was accomplished as shown in Scheme 3. Suzuki coupling between an appropriate boronic acid and commercially available 2-bromo-4-methyl imidazole afforded the corresponding 2-phenyl derivatives **17a**-**d** that were processed as described in Scheme 1 to afford the respective 2- and 4-chlorophenyl thiazolyl analogs **18b** and **18c**, respectively as well as the 3-methoxy and 3-trifluoromethoxy targets **18a** and **18d**. In both sets of analogs, we elected to use only the optimal 2-thiazolyl amide to provide the best comparison for activity versus **10a**-**h**.

**Scheme 2:**
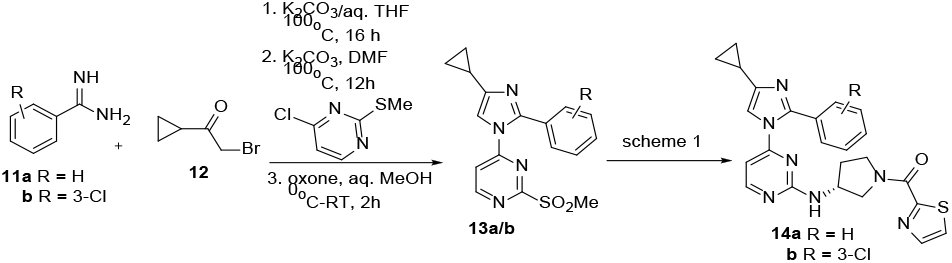
4-Cyclopropyl PfPKG Inhibitors **14a-b**

**Scheme 3:**
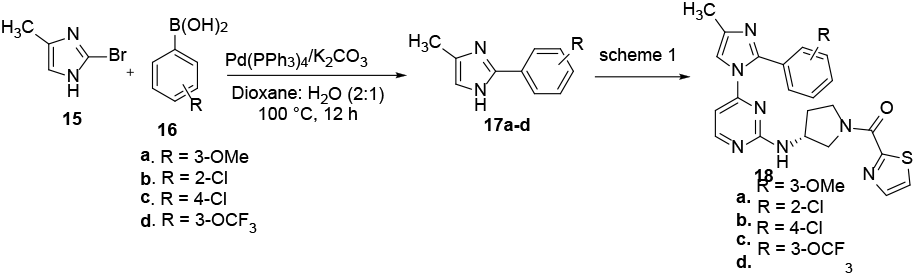
Phenyl-substituted PfPKG Inhibitors **18a-d**

Evaluation of these compounds as PfPKG inhibitors revealed that the cyclopropyl group in **14a** provides a three-fold improvement in potency compared to **10a** (Table 2) and that 3-chloro substitution provides a small additional benefit in **14b**. Among the group of phenyl substituents evaluated, 3-chloro is preferred to its regioisomers **18b** and **18c**, similar to our previous observations. The other 3-substituted derivatives examined in this small set are comparable (**18a, 18d**) to **14a** or less potent (**18b, 18c**).

**Table 2:** *in vitro* PfPKG inhibition **14a/b**-**18a-d**

| Cpd | IC<sub>50</sub><br>(nM)/%<br>inh @<br>1 μM |
| --- | --- |
| **14a** | 100 |
| **14b** | 60 |
| **18a** | 86 |
| **18b** | 47% |
| **18c** | 1160 |
| **18d** | 150 |

The most potent example in this set, **14b**, was examined for inhibition of human PKG and *P. falciparum* mutant T618Q, and as observed previously, demonstrated excellent selectivity against these two kinases with no inhibition at 10 μM. These results led us to examine **14a** and **14b** in more detail with a focus on cellular activity and *in vitro* ADME. These imidazole-based PfPKG inhibitors have moderate to good water solubility, and submicromolar inhibition of CYP3A4 (Table 2). Both compounds show poor metabolic stability and metabolite ID studies are underway to guide solutions to this important issue. The positive data in Table 3 shows neither **14a** nor **14b** have measurable hERG activity, unlike the thiazole example in Figure 1.

**Table 3:** *in vitro* ADME Characterization **14a-b**

| Cpd | HLM<br>t<sub>1/2</sub><br>(min) | MLM<br>t<sub>1/2</sub><br>(min) | H<sub>2</sub>O<br>sol<br>(μM) | 3A4<br>IC<sub>50</sub><br>(μM) | 2C9<br>IC<sub>50</sub><br>(μM) | 2D6<br>IC<sub>50</sub><br>(μM) | hERG<br>%<br>inh.<br>@ 10<br>μM |
| --- | --- | --- | --- | --- | --- | --- | --- |
| **14a** | 4.8 | < 2 | 92 | 0.29 | > 10 | > 10 | 1 |
| **14b** | 3.7 | < 2 | 28 | 0.19 | 2.8 | > 10 | 9 |

Using the HepG2-*P. berghei* Luc (PbLuc) sporozoite infectivity assay and **5** as a positive control, we chose to evaluate **3, 4, 10a** and **14b** to investigate a correlation between enzymatic and cellular activity. We were encouraged by the strong activity displayed the imidazoles (Table 4). The more efficacious imidazole, **14b**, like **5**, does not show a dose response (>90% at 2 and 10 μM) and exhibits comparable activity to **5**. Although it is not definite from the limited concentrations employed in this assay, **14b** appears to be more efficacious compared to the **10a** in this initial screen, suggesting a potential correlation between *in vitro* enzymatic inhibition and cellular efficacy. This positive data contrasts with the poor activity of isoxazoles **3** and **4**.

**Table 4:** *P. berghei* sporozoite screening infectivity assay

| Cpd | % decr<br>infn ±<br>SD @ 2<br>μM | % decr<br>infn ±<br>SD @<br>10 μM | P value |
| --- | --- | --- | --- |
| **3** | 52±15 | 55±21 | >0.05 |
| **4** | 0 | 22±38 | >0.05 |
| **10a** | 78±20 | 89±7 | < 0.001 |
| **14b** | 94±3 | 93±3 | < 0.001 |
| **5** | 91±8 | 94±4 | < 0.001 |

A more quantitative examination of **14a** and **14b** against asexual blood stages (3D7) and PbLuc sporozoite-HepG2 infectivity revealed distinct differences between **5, 14a** and **14b** (Table 5). This data shows that **5** is more effective in these cellular assays compared to imidazoles **14a** and **14b**. We note that the approximate 20-fold EC_50_ difference in the PbLuc HepG2 assay between the more efficacious **5** and **14b** supports the observation above of a correlation between *in vitro* PfPKG potency and cellular activity in the imidazole series. The modest activity exhibited by **14a** and **14b** in the asexual blood stage assay is not surprising since the asynchronous asexual replication of *P. falciparum* makes PfPKG inhibition less effective at this stage of the life cycle.^8^

**Table 5:** *P. falciparum* blood stage & *P. berghei* sporozoite EC_50_ values

| Cpd | EC<sub>50</sub> (μM)<br>*P. falciparum* 3D7 | EC<sub>50</sub> (μM)<br>*PbLuc* hepatic stage |
| --- | --- | --- |
| **5** | 0.15 | 0.3 |
| **14a** | 7.6 | 20 |
| **14b** | 3.3 | 5.7 |

This encouraging data led us to examine these imidazoles against additional *Plasmodium* species in other cell-based assays to more completely characterize this series and provide a baseline for future work. Imidazoles **10a** and **14b** were investigated in a dose-response assay that evaluated *P. cynomolgi* infectivity in prophylactic and radical cure modes. These two examples were selected to provide additional evidence to support the hypothesis of a correlation between *in vitro* PfPKG potency and cellular efficacy against other *Plasmodium* species. The prophylactic assay is a measure of the ability of *P. cynomolgi* to form hepatic schizonts or hypnozoites. The radical cure mode evaluates activity against existing schizonts and hypnozoites. The data in Table 6 show that **14b** is more active than **10a** in the prophylactic mode with a sub-micromolar IC_50_, and interestingly has a modest effect in the radical cure mode against schizonts, comparable to tafenoquine. This data is consistent with a dose response effect for both compounds and indicates **14b** is more efficacious than **10a** in cell-based anti-parasitic assays. Efficacy in the prophylactic schizont model is important because it demonstrates potential for interrupting the parasite’s liver development post sporozoite invasion of hepatocytes. Reducing liver infection by human-infective sporozoites is known to reduce severity and incidence of malaria.^6^ The data in Table 6also show these two compounds, unlike the controls, are comparatively non-toxic to host cells.

**Table 6:** *P. cynomolgi* pre-erythrocytic stage assays

| Compound | Prophylactic Mode |  |  |
| --- | --- | --- | --- |
|  | Schizont<br>IC<sub>50</sub><br>(μM) | Hypnozoite<br>IC<sub>50</sub><br>(μM) | Toxicity<br>(μM) |
| **14b** | 0.93 | 9.88 | >20 |
| **10a** | 7.91 | > 20 | >20 |
| Maduramicin | 0.01 | 0.02 | 5.78 |
| Tafenoquine | 0.19 | 0.14 | 13.82 |
| Compound | Radical Cure Mode |  |  |
|  | Schizont<br>IC<sub>50</sub><br>(μM) | Hypnozoite<br>IC<sub>50</sub><br>(μM) | Toxicity<br>(μM) |
| **14b** | 4.62 | > 20 | > 20 |
| **10a** | 11.59 | > 20 | > 20 |
| Maduramicin | 0.03 | 0.03 | 11.91 |
| Tafenoquine | 3.61 | 3.50 | 18.61 |

When comparing these new PfPKG inhibitors to known inhibitors (Figure 3), it is not surprising that enzymatic potency, while important, is not the sole determinant for cellular activity. It is encouraging to note the cellular activity of imidazoles **14a** and **14b**, in contrast to the more potent isoxazoles. One observation based on data in this paper is that molecular weight contributes to cellular activity; a comparison of **3, 4, 14b** and **5** shows that the two highest molecular weight compounds show at best weak activity in cells. There does not appear to be a direct relationship between cLogP and cellular activity (cf. **5, 4** and **14b**).

**Figure 3:**
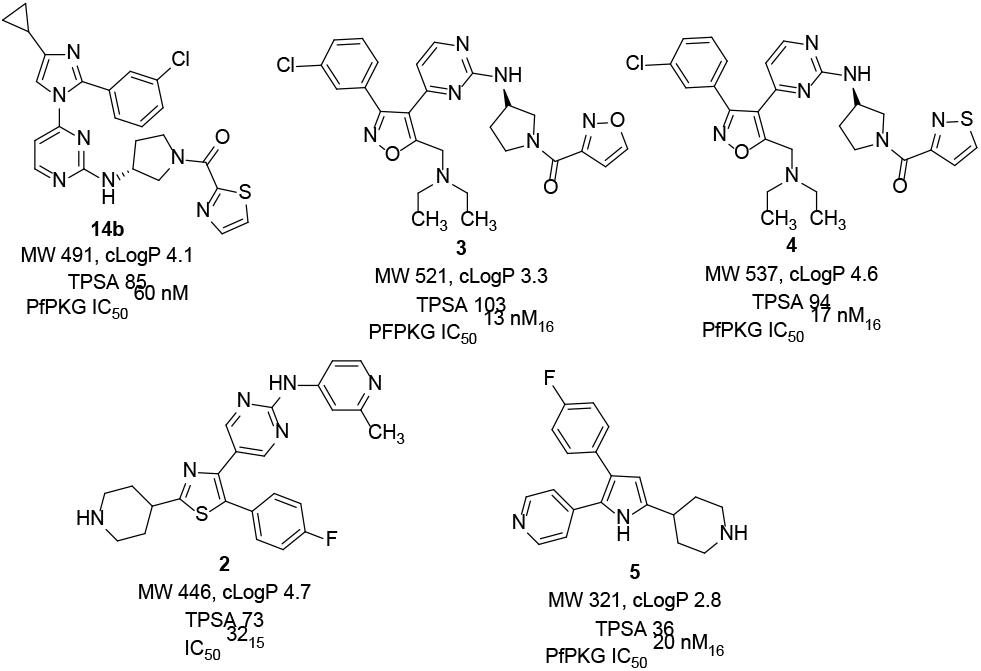
Chemical Property Comparison

We have shown that **5** and the two isoxazoles are competitive PfPKG inhibitors that bind in the ATP pocket of the enzyme^19^. Studies are ongoing with imidazoles such as **14b** to determine to mode of action for this class of compounds, along with investigations into obtaining what would be the initial report of an inhibitor bound to *P. falciparum* PKG.

In conclusion, in response to the need to identify novel chemotypes as PfPKG inhibitors that lack the safety issues associated with many known scaffolds, we report the discovery and initial characterization of a new imidazole-based chemotype with good *in vitro* PfPKG inhibition, and promising cellular activity that includes a correlation between *in vitro* enzymatic activity and efficacy, lacks the hERG issues associated with other chemotypes and does not have any structural alerts associated with genotoxicity. Initial structure-activity relationships are distinct from known PfPKG inhibitors and while the ADME profile of lead **14b** has weaknesses that are not unusual in early leads, the positive aspects of the imidazole series provide the impetus to address the pharmaceutical property issues that currently exist. Those efforts are ongoing and will be reported in due course.

## ASSOCIATED CONTENT

### Supporting Information

Full experimental details on the synthesis and characterization of compounds, *in vitro* enzyme assay, cellular parasite infectivity and *in vitro* ADME assays are provided along with the manuscript in review cited as reference 19.

The Supporting Information is available free of charge on the ACS Publications website.

Chemistry-synthesis and characterization PDF

Biology: *in vitro* PfPKG assays, cellular parasite infectivity, *in vitro* ADME PDF

## Author Contributions

All authors have given approval to the final version of the manuscript.

## Funding Sources

This research was supported by the Sokol Institute for Pharmaceutical Life Sciences (JJS and DPR), by NIH RO1-AI-133633-01 (JJS, PB and DPR) and by the Military Infectious Disease Research Program Q0480_19_WR_CS_OC for BSP and PJL

## ACKNOWLEDGMENT

We acknowledge the Entomology Branch and Veterinary Medicine Branch AFRIMS, with special thanks to Ratawan Ubalee and team for the production of *P. cynomolgi*-infected mosquitoes.

## ABBREVIATIONS

PfPKG: *Plasmodium falciparum*
cGMP: dependent protein kinase
PbLuc: *Plasmodium berghei* luciferase
SAR: structure-activity relationship
ATP: adenosine triphosphate
ADME: absorption, distribution, metabolism, elimination
CYP3A4: cytochrome P450 3A4
hERG: human ether-a-go-go related gene.

---

Authors are required to submit a graphic entry for the Table of Contents (TOC) that, in conjunction with the manuscript title, should give the reader a representative idea of one of the following: A key structure, reaction, equation, concept, or theorem, etc., that is discussed in the manuscript. Consult the journal’s Instructions for Authors for TOC graphic specifications.

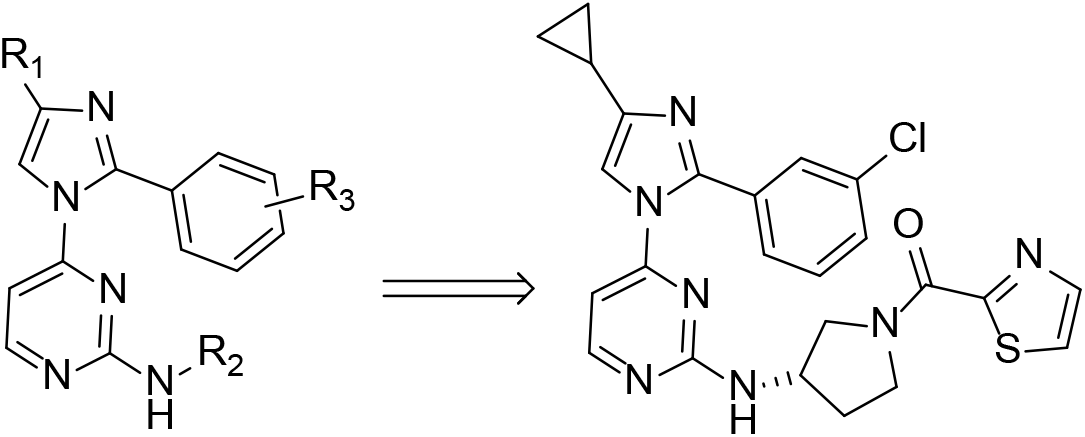

## Notes

### Competing Interest Statement

The authors have declared no competing interest.

