## Supplemental information for "Discovery of imidazole-based inhibitors of *P. falciparum* cGMP-dependent protein kinase"

#### Discovery of novel potent, cell-permeable inhibitors of *P. falciparum* cGMP-dependent protein kinase

+Sokol Institute for Pharmaceutical Life Sciences and Department of Chemistry and Biochemistry, Montclair State University, Montclair, New Jersey 07043, United States; %-Moulder Center for Drug Discovery Research, Temple University, Philadelphia PA, 19140; $-Department of Microbiology, Biochemistry and Molecular Genetics, Rutgers New Jersey Medical School, Newark NJ 07103; #-Department of Drug Discovery. Experimental Therapeutics Branch, Walter Reed Army Institute of Research, 503 Robert Grant Avenue, Silver Spring MD 20910.

***

##### General chemical procedures:

**Compound characterization:** All reagents and solvents were used as received from commercial suppliers. Compounds were analyzed using a UPLC system with a BEH-C18column (2.1 cm x 50 mm, 0-100% acetonitrile/water gradient over 6 minutes with UV 254 nmdetection) and an LCMS 2020 system (LC18 25 cm x 4.6 mm 5 μM column, 0-95%acetonitrile/water gradient over 10 minutes; UV monitor at 220 and 254 nM). Thin layer chromatography was done on silica gel G plates with UV or phosphomolybdic acid detection. Mass spectra were recorded on a CMS system with ESI probe. All of the reported yields are for isolated products and compounds were purified by automated flash chromatography. Proton NMR and $^{13}C$ spectra were obtained at 400 and 101 MHz, respectively, in $CDCl_3$ unless otherwise stated. All final compounds had purities of at least 95% based on 1H NMR and UPLC analyses.

##### General route for synthesis of 10a-h:

**Arylation:** To the solution of **7** (1 eq) in anhydrous DMF (10 ml) at ambient temperature was added NaH (1.2 eq, 60% in mineral oil) and the resulting mixture was stirred for 30 minutes. at 0°C followed by addition of 4-chloro-2-(methylthio)pyrimidine (1.2 eq). The resulting reaction mixture then was stirred for 12 h with temperature gradually rising from 0 °C to ambient temperature, then quenched by addition of water (10 ml). The quenched mixture was partitioned between ethyl acetate/$H_2O$ (3 times). Organic layer was collected, washed with brine, dried, concentrated and purified by CombiFlash silica gel chromatography eluting with a mixture of ethyl acetate/hexane to provide the desired product as an off-white solid.

**9:** Pyrimdinyl methyl sulfide was dissolved at room temperature in a mixture of methanol/distilled water (2:1 ratio, final concentration ~0.2M). Oxone (potassium peroxymonosulfate, 2.1 eq) was added at room temperature and the reaction was stirred overnight. The reaction mixture was poured into water and extracted with 4 portions of ethyl acetate. The collected organic extracts were washed with brine, dried, concentrated and purified by CombiFlash silica gel chromatography eluting with a mixture of ethyl acetate/hexane to provide the desired sulfone as a white solid.

**Amine displacement of sulfone**: **9** was dissolved in reagent grade DMSO (~0.3M) at room temperature. Triethylamine (3 equivalents) were added, followed by 2 equivalents of the desired Boc-protected diamine. The reaction was heated under nitrogen to 65 °C overnight. After cooling to room temperature the reaction mixture was poured into brine solution and extracted 4 times with ethyl acetate. The collected organic extracts were washed with brine, dried and concentrated. The desired product was isolated by CombiFlash silica gel chromatography eluting with 5-10% methanol in dichloromethane to furnish pure product.

**Boc Deprotection**: The respective Boc-amine (1 eq) was dissolved in 10 mL methylene chloride was added excess trifluoro acetic acid at 0 °C. The reaction mixture was stirred for 2 hours at room temperature. Solvent was removed in vacuo, the residue was dissolved in brine solution and quenched it with solid NaHCO3 till the pH is neutral and extracted 4 times with ethyl acetate. The collected organic extracts were washed with brine, dried and concentrated. The desired product was isolated by CombiFlash silica gel chromatography eluting with 10% methanol in dichloromethane to furnish pure product.

**Amide coupling**: To the solution of the appropriate amine in reagent grade DMF at room temperature was added 3 equivalents of pyridine, followed by 1.3 equivalents of HATU and 1.2 equivalents of the desired carboxylic acid. The reaction was stirred at room temperature overnight. In the morning, the reaction mixture was poured into 5 volumes of water and extracted with 4 portions of ethyl acetate. The collected organic extract was washed with two portions of brine, dried and concentrated. The product was purified by CombiFlash silica gel chromatography eluting with 5-10% methanol in dichloromethane to furnish the desired amine product.

Yields stated with each compound reflect this last reaction.

**10a** Yield: 62%; δ 8.13 (dd, *J* = 5.4, 1.6 Hz, 1H), 7.90 (dd, *J* = 7.4, 3.2 Hz, 1H), 7.54 (dd, *J* = 3.1, 1.4 Hz, 1H), 7.49 – 7.40 (m, 2H), 7.40 – 7.31 (m, 3H), 7.29 (d, *J* = 0.9 Hz, 1H), 7.10 – 6.72 (m, 5H), 6.17 (s, 1H), 5.50 (s, 1H, NH), 4.42 – 4.26 (m, 2H), 4.02 – 3.68 (m, 3H), 2.31 (s, 3H), 2.01 (s, 1H), 1.86 (dd, *J* = 14.5, 7.5 Hz, 1H); $^{13}$C δ 165.3, 165.2, 161.8, 159.7, 159.4, 159.1, 157.3, 146.5, 143.9, 143.8, 143.6, 143.6, 138.5, 129.0, 128.4, 124.2, 116.4, 116.3, 47.1, 45.7, 32.3, 29.5, 13.6. ESI-MS: m/z 432.0 [M + H]$^+$.

**10b** Yield: 58%; white solid; δ- 8.39 (dd, *J* = 9.8, 6.4, 1H), 8.12 (dd, *J* = 5.4, 0.9 Hz, 1H), 7.37 – 7.26 (m, 6H), 6.77 (d, *J* = 2.3 Hz, 1H), 6.12 (s, 1H), 4.17 (s, 1H), 4.06 (dd, *J* = 14.1, 7.2 Hz, 1H), 3.91 – 3.84 (m, 4H), 3.77 (dd, *J* = 19.6, 13.7 Hz, 2H), 2.28 (s, 3H), 2.12 (s, 1H), 1.94 (s, 1H). $^{13}$C: δ 162.2, 161.8, 157.0, 147.5, 147.4, 140.3, 137.6, 137.5, 135.2, 129.0, 128.6, 128.4, 128.3, 109.1, 108.7, 46.9, 44.8, 22.7, 13.1. ESI-MS: m/z 429.0 [M + H]$^+$.

**10c** Yield: 60%; white solid; δ 8.36 (dd, *J* = 10.4, 5.0 Hz, 1H), 8.15 – 7.78 (m, 1H), 7.91 – 7.87 (m, 1H), 7.54 – 7.26 (m, 6H), 6.49 (dd, *J* = 18.5, 4.5 Hz, 1H), 5.49 (s, 1H), 4.55 – 4.18 (m, 2H), 4.08 (dd, *J* = 12.5, 4.9 Hz, 1H), 3.67 (dd, *J* = 85.2, 36.0 Hz, 2H), 2.31 (s, 3H), 2.18 – 1.94 (m, 1H), 1.94 – 1.65 (m, 1H). $^{13}$C: δ 165.2, 162.1, 160.6,

160.3, 159.1, 134.2, 128.5, 128.3, 124.2, 124.1, 114.9, 114.8, 47.1, 45.7, 38.6, 12.0. ESI-MS: m/z 432.0 $[M + H]^+$.

**10d** Yield: 53%; white solid; δ 8.35 (s, 1H), 7.89 (s, 1H), 7.68 (s, 1H), 7.54 (s, 1H), 7.47 – 7.31 (m, 5H), 6.47 (s, 1H), 6.22 (s, 1H), 4.19 (s, 2H), 4.06 – 3.73 (m, 1H), 3.48 – 3.36 (m, 2H), 2.33 (s, 3H), 1.74 (s, 2H), 1.59 (s, 2H); $^{13}$C: δ 162.2, 160.3, 143.1, 142.7, 132.0, 129.1, 128.6, 128.4, 128.3, 127.6, 127.4, 124.1, 123.7, 50.5, 47.7, 44.2, 34.7, 31.6, 25.3, 22.6, 11.9; ESI-MS: m/z 446.0 $[M + H]^+$.

**10e** Yield: 57%; off-white solid; δ-8.37 (dd, *J* = 4.6, 2.6 Hz, 1H), 7.88 – 7.85 (m, 1H), 7.52 – 7.51 (m, 1H), 7.36 – 7.26 (m, 6H), 6.55 (s, 1H), 5.43 – 5.26 (m, 2H), 4.49 (s, 1H), 3.65 (s, 1H), 3.18 (s, 1H), 2.81 – 2.79 (m, 1H), 2.30 (s, 3H), 1.73 – 1.68 (m, 2H), 1.25 – 1.24 (m, 2H); $^{13}$C NMR: δ 165.1, 161.9, 161.8, 160.5, 160.3, 159.2, 143.2, 142.9, 134.3, 128.2, 124.1, 114.6, 48.3, 45.1, 42.6, 12.1; ESI-MS: m/z 446.0 $[M + H]^+$.

**10f:** Yield 68%, white solid; *δ*= 8.3(d, 1H, H11, J=4.9Hz), 7.9(s, 1H, H21), 7.5(s, 1H, H20), 7.4-7.1(m, 7H, H2, H5, H6, H7, H5', H6', H10), 6.4(s, 1H, NH), 4.5-3.4(m, 5H, H13,H14, H15), 2.3 (s, 3H, H8), 2.3-1.6 (m, 4H, H15, H16). ESI $(M+ H)^+$ +$(MeOH + H)^+$ 480 m/z

**10g**: Yield 61%; white solid; *δ*-8.3(d, 1H, H11, *J*=4.8Hz), 7.8(s, 1H, H20), 7.7 (s, 1H, NH), 7.6(s, 1H, H19), 7.4-7.2(m,6H, H2, H5, H6, H7, H5', H6'), 6.6 (s, 1H, NH), 6.4 (s,1H, H10), 3.4(s,2H, H13), 2.3 (s, 3H, H8), 1.0-0.8(m, 4H, H15,H16). ESI $(M+ H)^+$ +$(MeOH + H)^+$ 466 m/z

**10h**: Yield 71%; white solid;*δ*-8.3(d, 1H, H11, *J*=4.8Hz), 7.8(s, 1H, H20), 7.7 (s, 1H, NH), 7.6(s, 1H, H19), 7.4-7.2(m,6H, H2, H5, H6, H7, H5', H6'), 6.6 (s, 1H, NH), 6.4 (s,1H, H10), 3.4(s,2H, H13), 2.3 (s, 3H, H8), 1.0-0.8(m, 4H, H15,H16). ESI $(M+ H)^+$ +$(MeOH + H)^+$ 466 m/z

**Synthesis of 14a/b**

**Preparation of cyclopropyl imidazole derivatives:** A mixture of benzamidine (1 eq), 2-bromo-1-cyclopropylethanone (1.2 eq) and potassium carbonate (2 eq) in 2:1 ratio of THF and water (50 ml) was stirred at reflux condition for 18 hours. After cooling to room temperature, the reaction mixture was filtered off and evaporated in vacuo. Then the residue was poured into water, extracted with ethyl acetate, dried over sodium sulfate, and evaporated in vacuo. The crude product was purified by CombiFlash silica gel chromatography eluting with 30-50% ethyl acetate in hexane to furnish the desired products.

The synthesis of **13a/b**, amine displacement, deprotection and acylation was carried out as described above. The yield stated below is for the acylation reaction to afford the target compound.

**14a.** Yield: 73%; off-white solid; δ 8.19 (d, *J* = 4.6 Hz, 1H), 7.95 (dd, *J* = 11.4, 3.2 Hz, 1H), 7.80 (d, *J* = 2.7 Hz, 1H), 7.41 – 7.34 (m, 6H), 6.44 (s, 1H), 4.16 (d, *J* = 5.6 Hz, 2H), 3.74 – 3.39 (m, 3H), 1.98 (s, 1H), 1.88 (dd, *J* = 9.6, 5.1 Hz, 2H), 0.89 – 0.75 (m, 4H); $^{13}$C: δ 165.4, 165.3, 161.7, 159.2, 159.1, 157.15, 144.5, 144.4, 129.2, 128.5, 124.2, 114.8, 114.6, 47.1, 45.6, 32.4, 29.6, 8.7, 7.2; ESI-MS: m/z 458.0 $[M + H]^+$.

**14b.** Yield: 69%; off-white solid; δ 8.16 (d, *J* = 5.4, 2.4 Hz, 1H), 7.87 (dd, *J* = 4.9, 3.2 Hz, 1H), 7.52 – 7.48 (m, 2H), 7.41 – 7.09 (m, 4H), 6.18 (s, 1H), 4.29 (d, *J* = 5.3 Hz, 2H), 4.0 - 3.63 (m, 3H), 2.16 (s, 1H), 1.91 – 1.85 (m, 2H), 0.89 – 0.83 (m, 4H); $^{13}$C

NMR ($CDCl_3$, 101 MHz): δ 165.3, 161.8, 159.1, 157.0, 145.0, 143.6, 134.3, 129.1, 124.2, 115.3, 115.1, 47.1, 45.6, 32.4, 29.6, 8.8, 7.1; ESI-MS: m/z 492.0 $[M + H]^+$.

**Synthesis of 18a-d:**

**17a-d:** A solution of 2 M $K_2CO_3$ solution (3 ml) in dioxane (3 ml) was taken in round bottom flask and was purged with nitrogen balloon for 5 minutes at room temperature. A mixture of aryl boronic acid (**16a-d**, 1.2 eq) and 2-bromo 4-methyl imidazole (**15**, 1 eq) was added to this reaction mixture, and it was again purged with nitrogen for 5 min. $Pd(PPh_3)_4$ (0.05 eq) was then added, followed by purging and allowed the reaction mixture to stir at 90 °C for overnight. After completion of the reaction, product was extracted with ethyl acetate (2 × 50 ml). The combined organic layer was concentrated in vacuo and crude reaction mixture was purified by CombiFlash silica gel chromatography eluting with 30-50% ethyl acetate in hexane to furnish the desired products.

**17a-d** were processed as described above to obtain **18a-d**. The yield for each compound is for the acylation reaction to obtain the target compound.

**18a.** Yield: 71%; off-white solid; δ 8.10 (dd, *J* = 5.4, 2.7 Hz, 1H), 7.87 (dd, *J* = 7.4, 3.2 Hz, 1H), 7.51 (dd, *J* = 3.1, 2.6 Hz, 1H), 7.28 – 7.20 (m, 2H), 7.04 – 7.01 (m, 1H), 6.94 – 6.88 (m, 2H), 6.13 (s, 1H), 4.41 (s, 1H), 4.23 (s, 1H), 3.93 – 3.83 (m, 1H), 3.76 – 3.74 (m, 4H), 3.54 (dd, *J* = 4.8, 4.0 Hz, 1H), 2.28 (s, 3H), 2.04 (dd, *J* = 6.2, 5.6 Hz, 1H), 2.10 – 1.78 (m, 1H); $^{13}C$ δ 165.3, 165.2, 161.8, 159.5, 157.0, 146.1, 143.6, 137.9, 129.5, 124.2, 124.1, 121.7, 121.4, 116.5, 116.4, 47.2, 45.7, 32.3, 29.5, 13.3; ESI-MS: m/z 462.0 $[M + H]^+$.

**18b.** Yield: 71%; off-white solid; δ 8.17 (d, *J* = 5.5 Hz, 1H), 7.89 (dd, *J* = 10.4, 3.2 Hz, 1H), 7.54 – 7.53 (m, 2H), 7.37 (dd, *J* = 12.8, 6.0 Hz, 4H), 6.28 (s, 1H), 5.85 (s, 1H), 5.29 (s, 1H), 4.21 (s, 2H), 3.80 – 3.51 (m, 3H), 2.37 (s, 3H), 2.06 – 2.03 (m, 1H), 1.80 (s, 1H); $^{13}C$ δ 165.2, 161.2, 159.1, 156.5, 143.9, 143.6, 133.9, 129.6, 127.1, 124.2, 115.1, 47.1, 45.6, 32.3, 29.5, 13.1; ESI-MS: m/z 466.0 $[M + H]^+$.

**18c.** Yield: 71%; off-white solid; δ 8.25 (s, 1H), 7.92 – 7.89 (m, 1H), 7.55 – 7.46 (m, 3H), 7.40 – 7.29 (m, 3H), 6.26 (s, 1H), 4.29 (s, 3H), 3.86 – 3.80 (m, 3H), 2.39 (s, 3H), 2.07 (s, 1H), 1.85 (s, 1H); $^{13}C$ δ 165.3, 161.8, 160.0, 159.2, 156.8, 143.7, 131.3, 131.2, 124.2, 116.8, 116.0, 47.1, 45.6, 32.3, 29.5, 12.8; ESI-MS: m/z 466.0 $[M + H]^+$.

**18d.** Yield: 71%; off-white solid; δ 8.27 (d, *J* = 5.3, 3.1 Hz, 1H), 7.92 – 7.88 (m, 2H), 7.56 – 7.53 (m, 2H), 7.48– 7.46 (m, 1H), 7.36 – 7.31 (m, 2H), 6.29 (s, 1H), 4.26 (s, 2H), 4.08 (s, 1H), 3.93 – 3.59 (m, 2H), 2.38 (s, 3H), 2.08 (s, 1H), 1.95 (s, 1H); $^{13}C$ δ 161.7, 159.7, 159.2, 157.4, 153.2, 149.9, 138.1, 129.0, 128.4, 119.7, 116.4, 46.8, 45.2, 32.3, 29.7, 13.3; ESI-MS: m/z 516.0 $[M + H]^+$.

#### $^1H$, $^{13}C$ and Mass spectra of final compounds

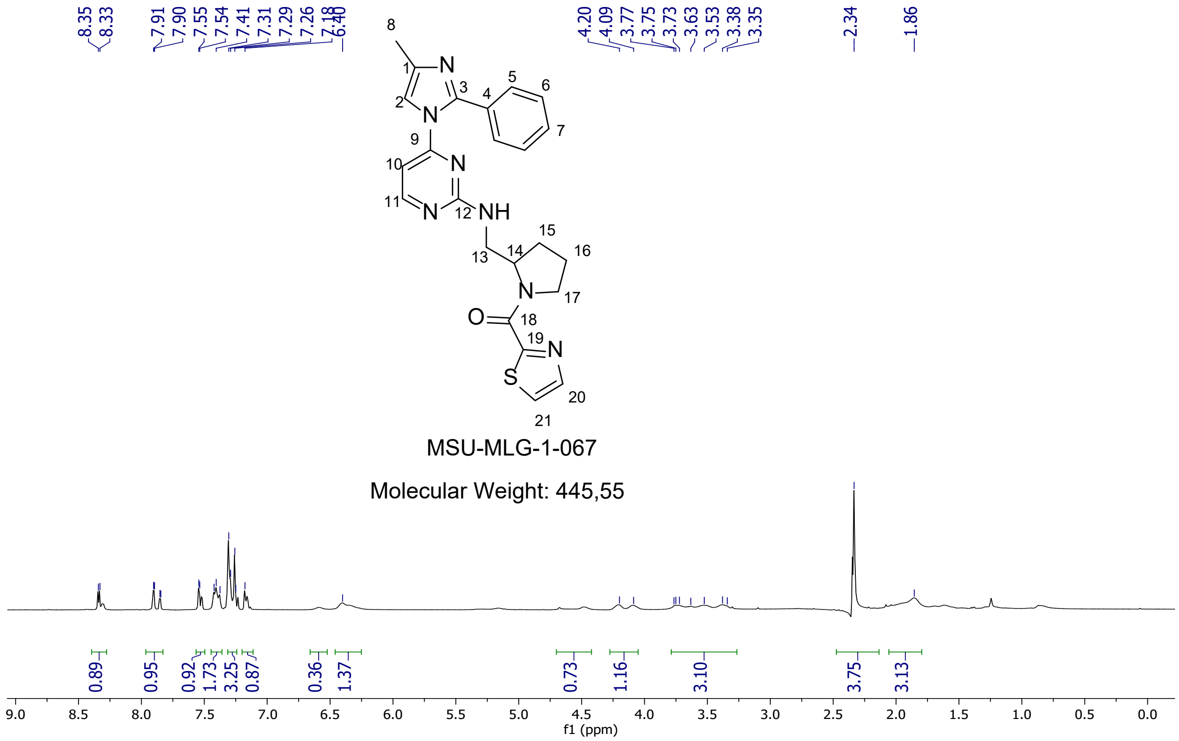


$^{1}H$ NMR ($CDCl_3$, 400Hz) 10f

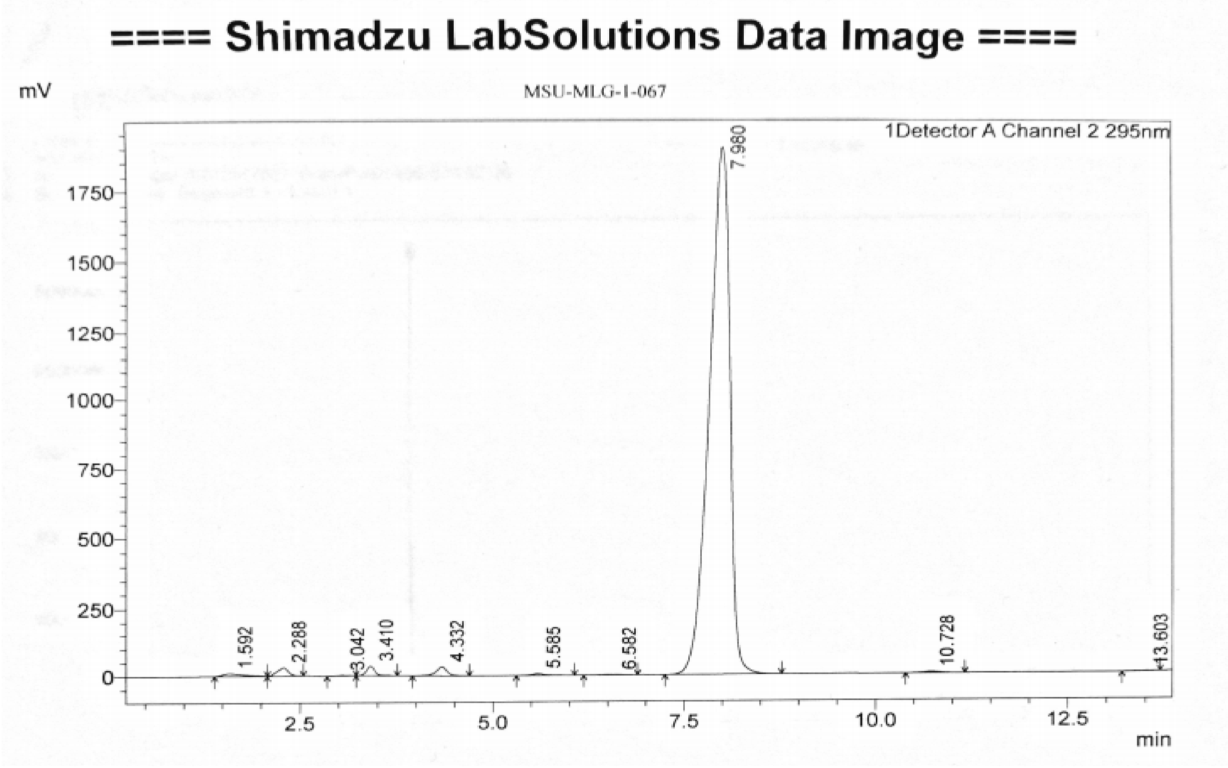


LC Spectrum 10f

**MS Spectrum** 10f $(M+H)^{+}$ +$(MeOH + H)^{+}$ 480 m/z

**10g**

**$^1$H NMR (400MHz, CDCl$_3$)**: $\delta$= 8.3(d, 1H, H11, *J*=4.8Hz), 7.8(s, 1H, H20), 7.7 (s, 1H, NH), 7.6(s, 1H, H19), 7.4-7.2(m,6H, H2, H5, H6, H7, H5', H6'), 6.6 (s, 1H, NH), 6.4 (s,1H, H10), 3.4(s,2H, H13), 2.3 (s, 3H, H8), 1.0-0.8(m, 4H, H15,H16) ppm. **ME** (M+ H)$^+$ +(MeOH + H)$^+$ 466 m/z

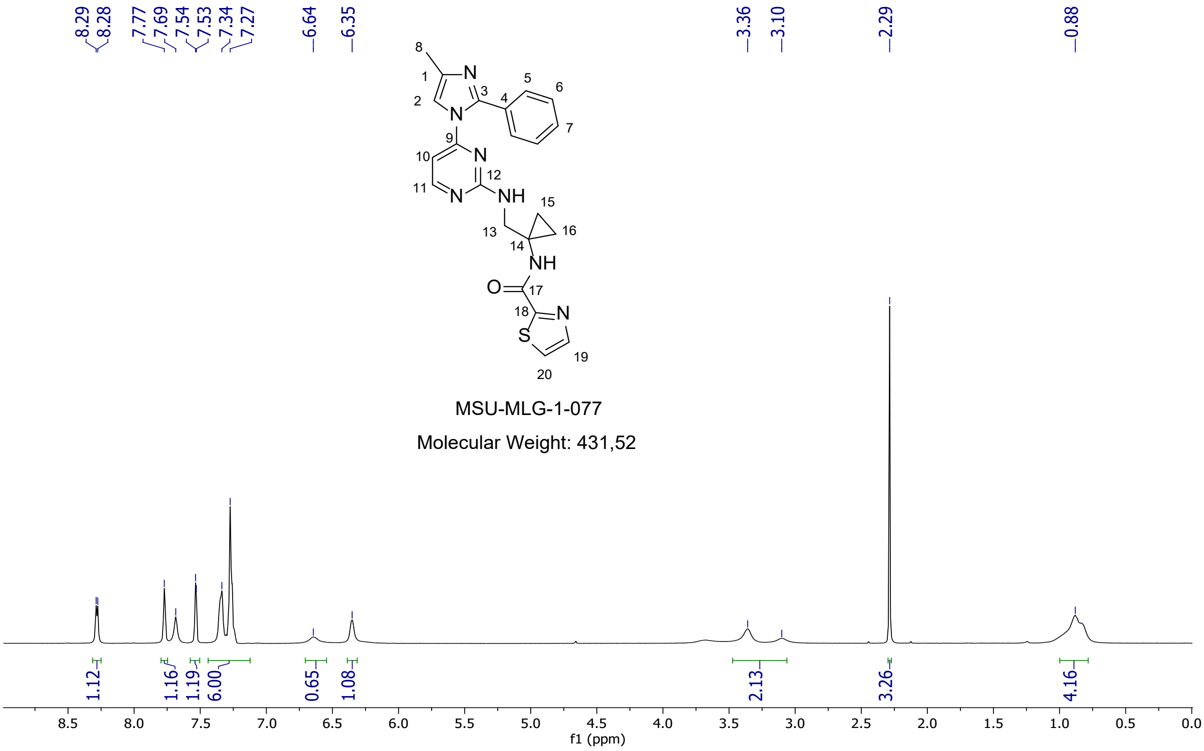


$^1$H NMR (CDCl$_3$, 400Hz) **10g**

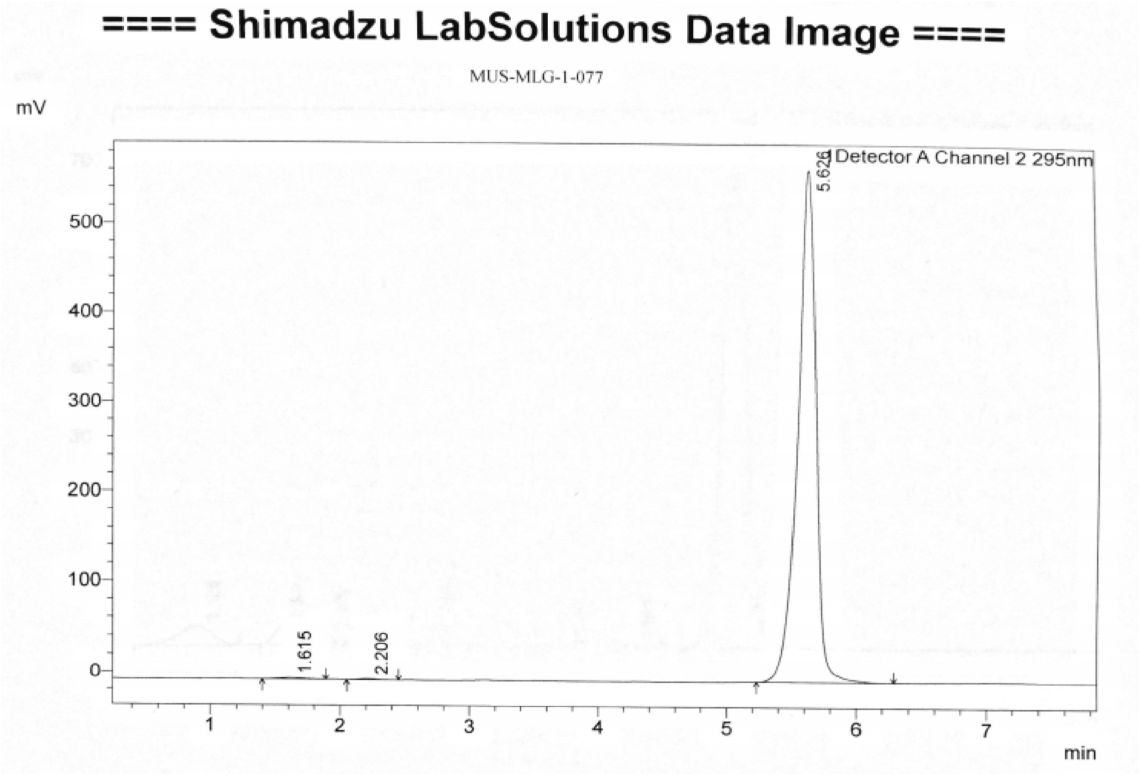


LC Spectrum **10g**

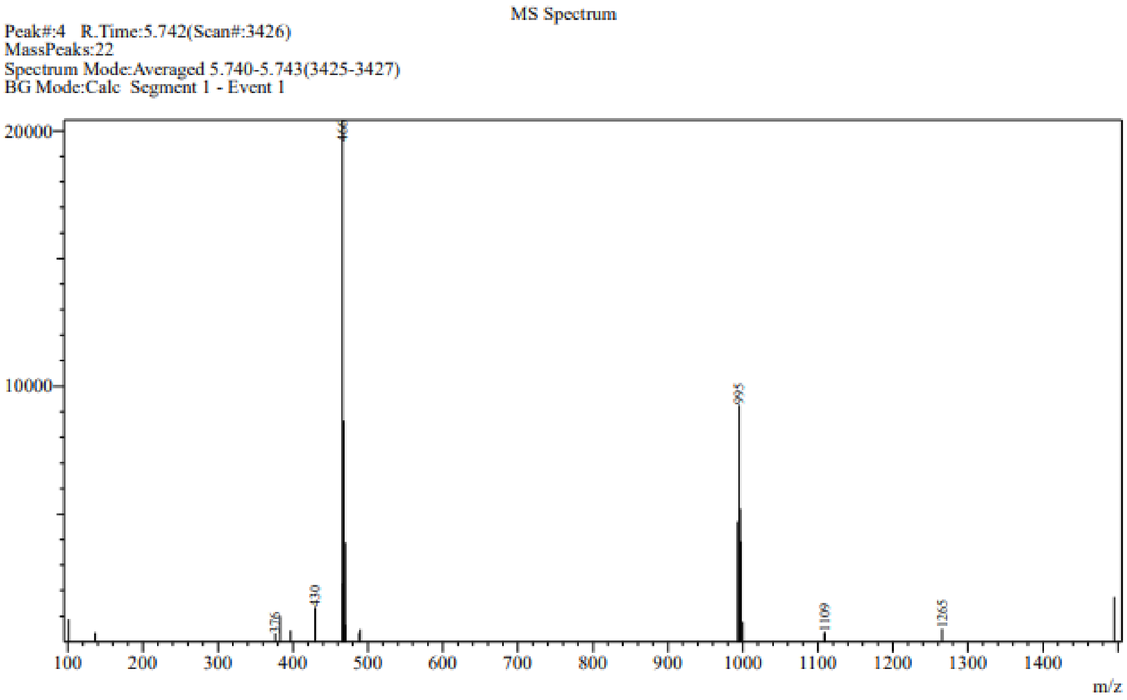


**MS Spectrum 10g** $(M+H)^+$ +$(MeOH + H)^+$ 466 m/z

**10h**

$^1$H NMR (400MHz, $CDCl_3$): δ= 8.3(d, 1H, H11, *J*=4.8Hz), 7.8(s, 1H, H20), 7.7 (s, 1H, NH), 7.6(s, 1H, H19), 7.4-7.2(m,6H, H2, H5, H6, H7, H5', H6'), 6.6 (s, 1H, NH), 6.4 (s,1H, H10), 3.4(s,2H, H13), 2.3 (s, 3H, H8), 1.0-0.8(m, 4H, H15,H16) ppm. **ME** $(M+H)^+$ $+(MeOH + H)^+$ 466 m/z

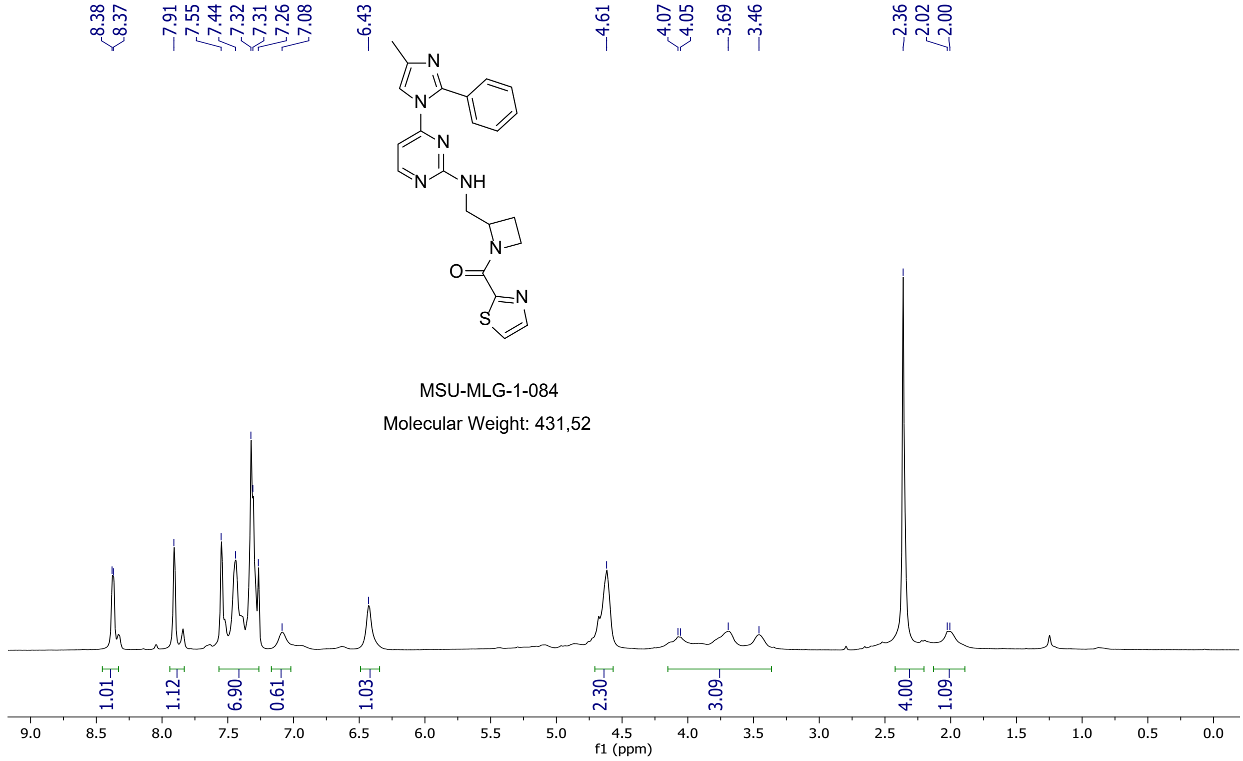


$^1$H NMR ($CDCl_3$, 400Hz) **10h**

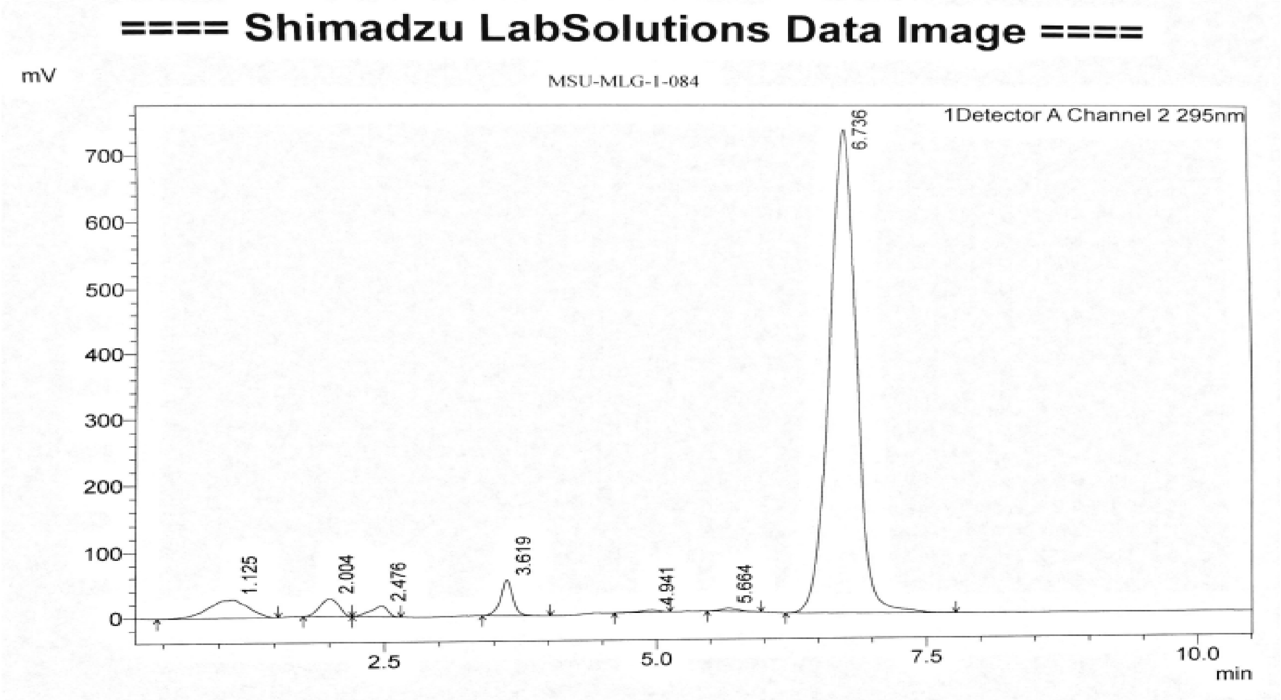

LC Spectrum **10h**

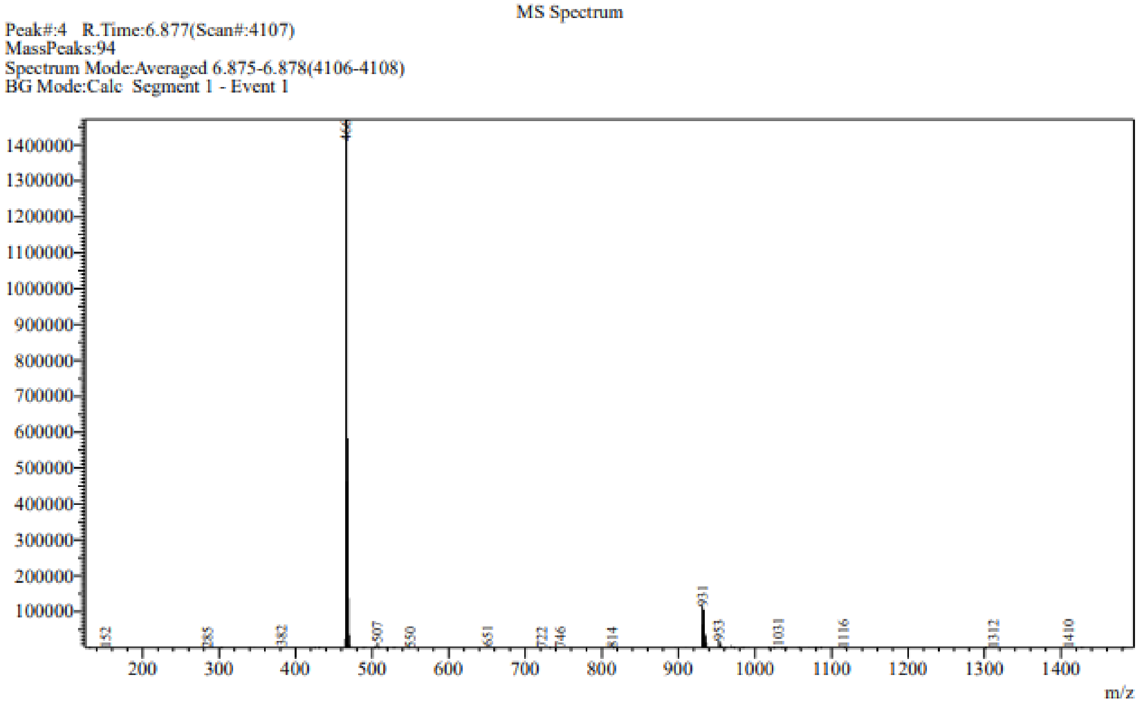


**MS Spectrum 10h** $(M+H)^+$ +$(MeOH + H)^+$ 466 m/z

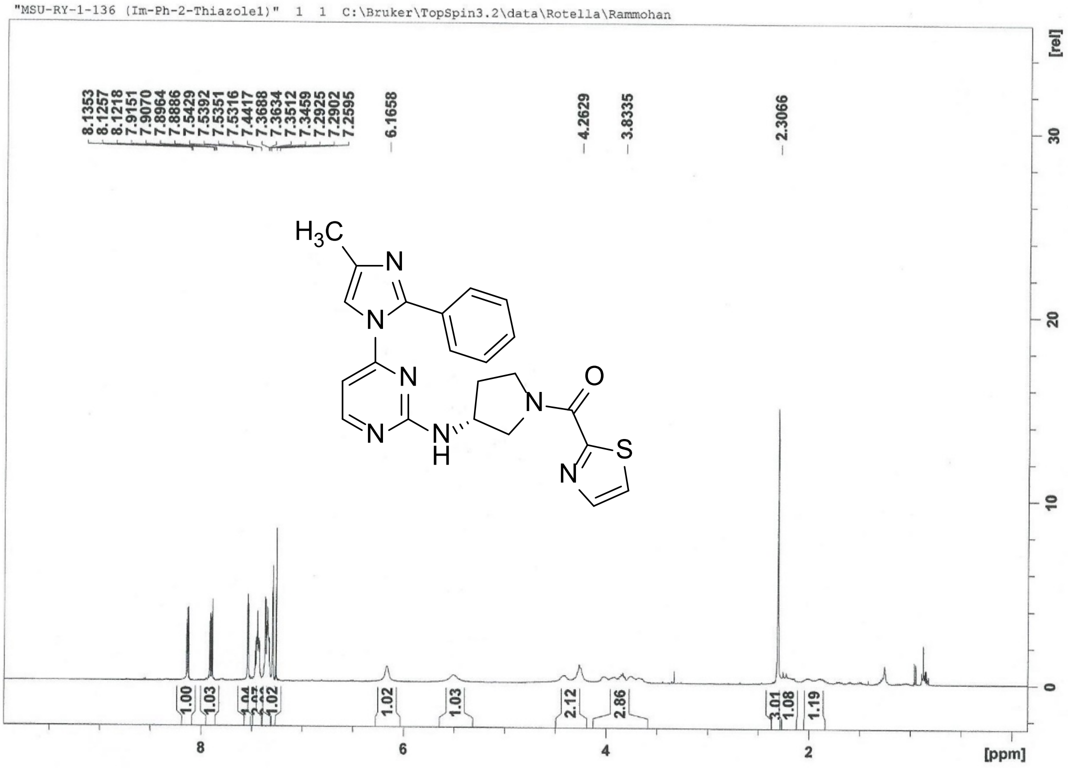
"MSU-RY-1-136 (Im-Ph-2-Thiazole1)" 1 1 C:\Bruker\TopSpin3.2\data\Rotella\Rammohan
8.1353
8.1257
8.1218
7.9151
7.9070
7.8964
7.8886
7.5429
7.5392
7.5351
7.5316
7.4417
7.3688
7.3634
7.3512
7.3459
7.2925
7.2902
7.2595
6.1658
4.2629
3.8335
2.3066
$H_3C$
N
N
N
N
NH
N
O
S
N
1.00
1.03
1.04
1.02
1.02
1.03
2.12
2.86
3.01
1.08
1.19
[rel]
30
20
10
0
8
6
4
2
[ppm]

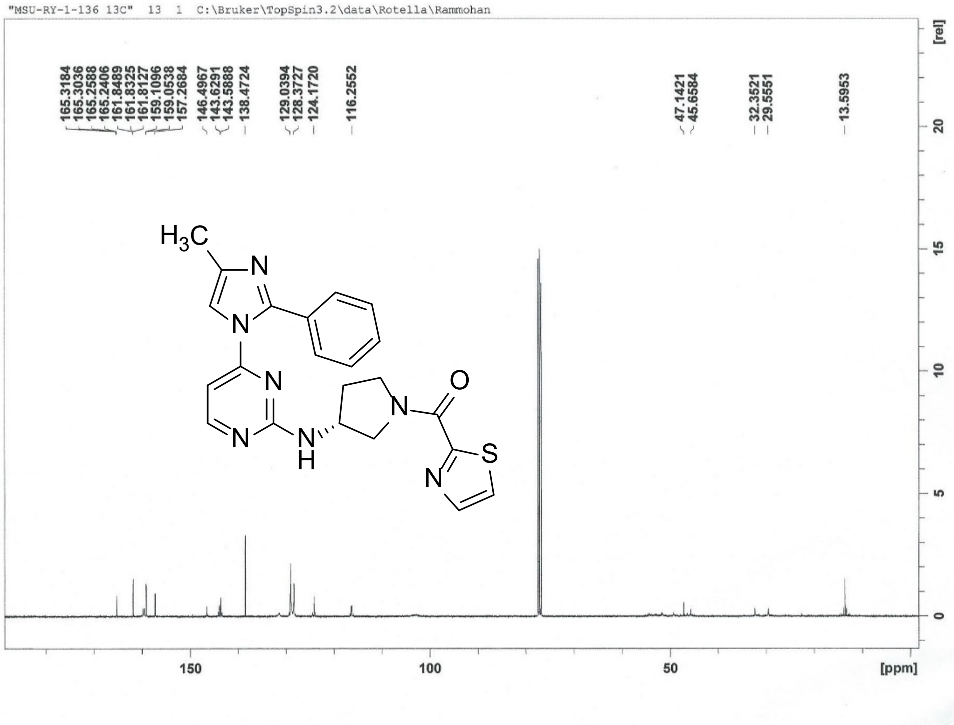
"MSU-RY-1-136 13C" 13 1 C:\Bruker\TopSpin3.2\data\Rotella\Rammohan
165.3184
165.3036
165.2588
165.2406
161.8489
161.8325
161.8127
159.1096
159.0538
157.2684
146.4967
143.6291
143.5888
138.4724
129.0394
128.3727
124.1720
116.2552
47.1421
45.6584
32.3521
29.5551
13.5953
$H_3C$
N
N
N
N
NH
N
O
S
N
[rel]
20
15
10
5
0
150
100
50
[ppm]

#### ==== Shimadzu LabSolutions Data Report ====

##### MSU-RY-1-136

Line#:1 R.Time:2.547(Scan#:----)
MassPeaks:35
RawMode:Averaged 2.545-2.548(1508-1510) BasePeak:432(52555)
BG Mode:Calc Segment 1 - Event 1

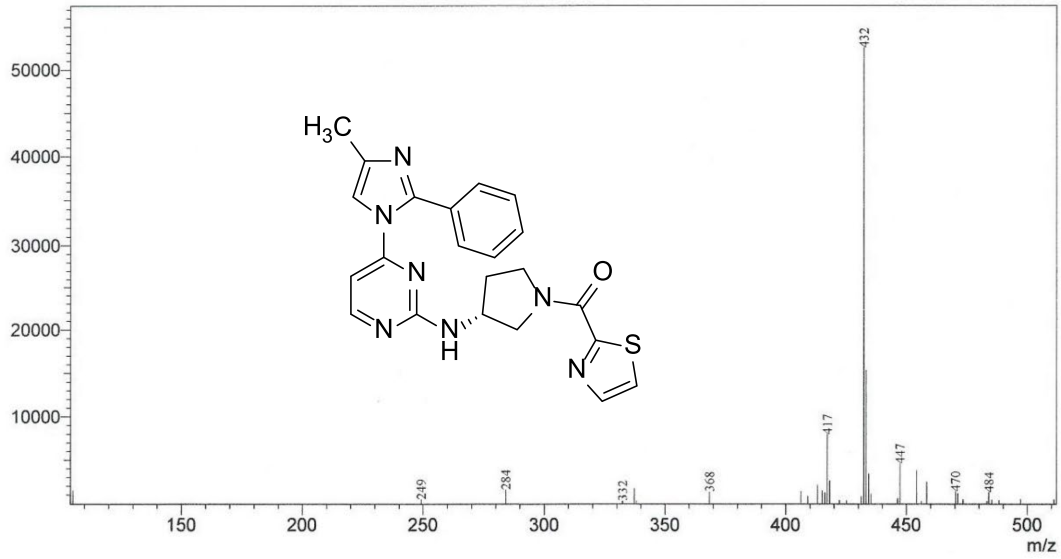


"MSU-RY-1-138 (Im-Ph-3-pyrazole)" 1 1 C:\Bruker\TopSpin3.2\data\Rotella\Rammohan

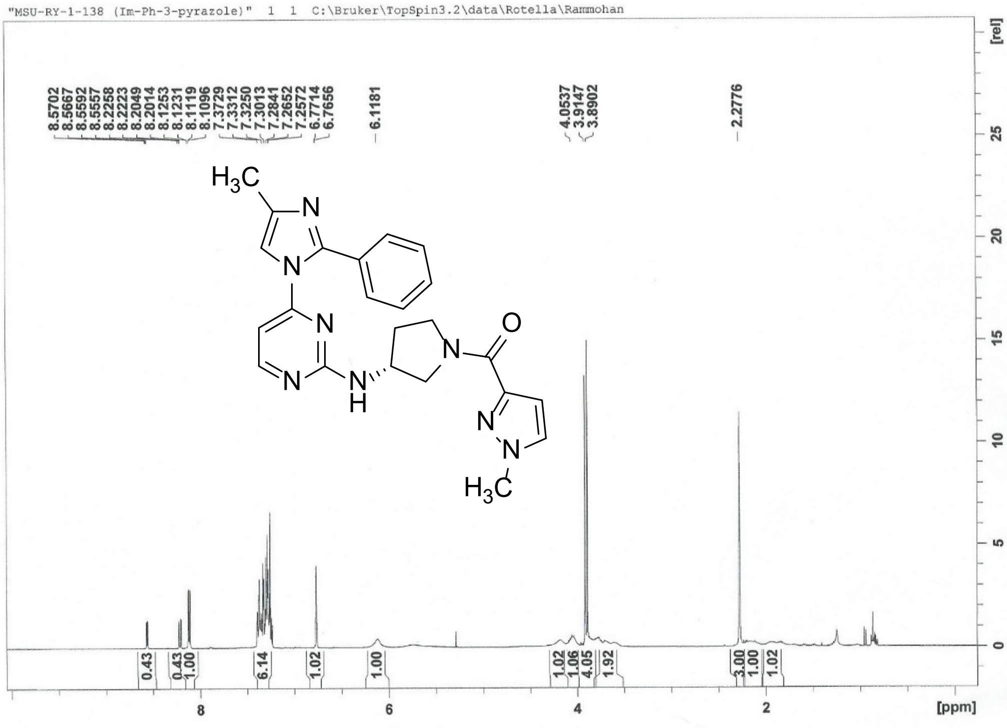

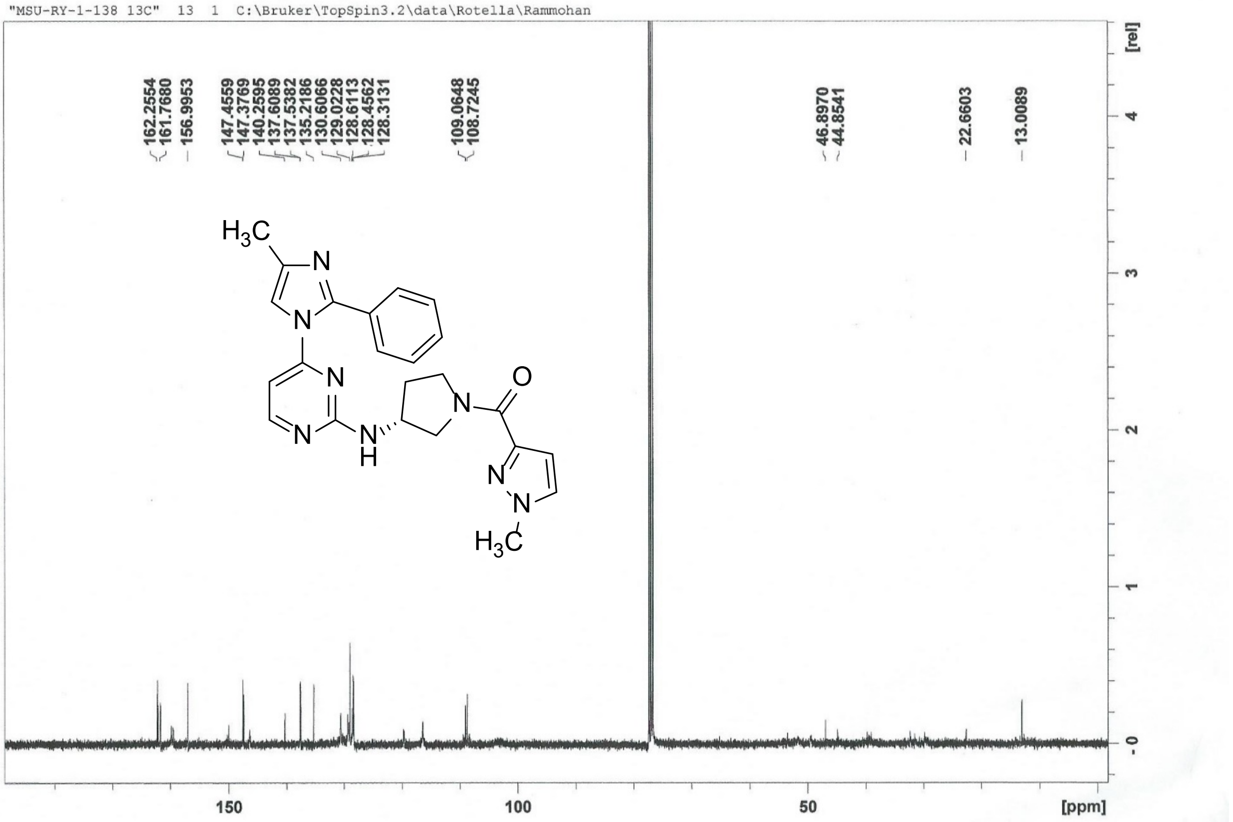
"MSU-RY-1-138 13C" 13 1 C:\Bruker\TopSpin3.2\data\Rotella\Rammohan
162.2554
161.7680
156.9953
147.4559
147.3769
140.2595
137.6089
137.5382
135.2186
130.6066
129.0228
128.6113
128.4562
128.3131
109.0648
108.7245
46.8970
44.8541
22.6603
13.0089
[rel]
150
100
50
[ppm]

### ==== Shimadzu LabSolutions Data Report ====

#### MSU-RY-1-138

Line#:1 R.Time:3.468(Scan#:----)
MassPeaks:20
RawMode:Averaged 3.467-3.470(2061-2063) BasePeak:429(303490)
BG Mode:Calc Segment 1 - Event 1

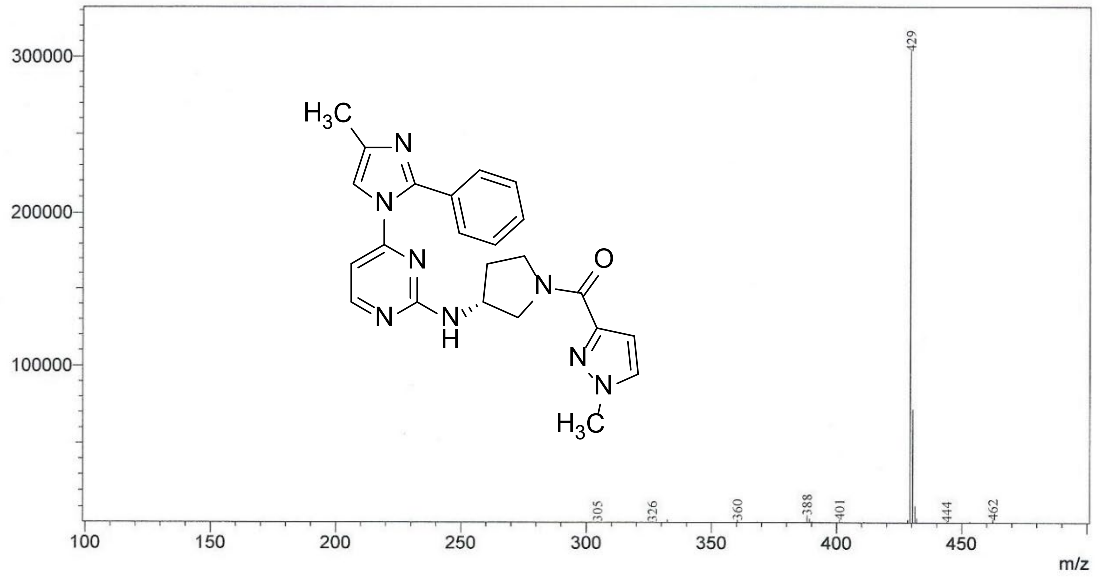

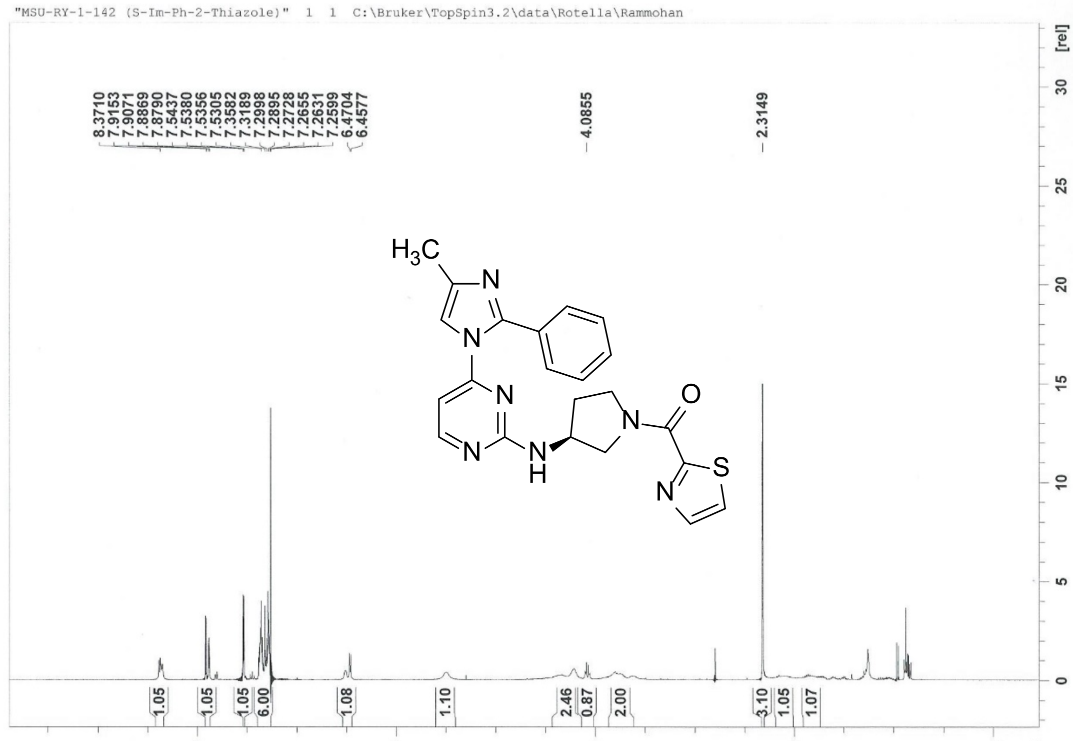
"MSU-RY-1-142 (S-Im-Ph-2-Thiazole)" 1 1 C:\Bruker\TopSpin3.2\data\Rotella\Rammohan
8.3710
7.9153
7.9071
7.8869
7.8790
7.5437
7.5380
7.5356
7.5305
7.3582
7.3189
7.2998
7.2895
7.2728
7.2655
7.2631
7.2599
6.4704
6.4577
4.0855
2.3149
H3C
N
N
N
N
N
H
N
O
S
N
1.05
1.05
1.05
6.00
1.08
1.10
2.46
0.87
2.00
3.10
1.05
1.07
[ppm]
[rel]

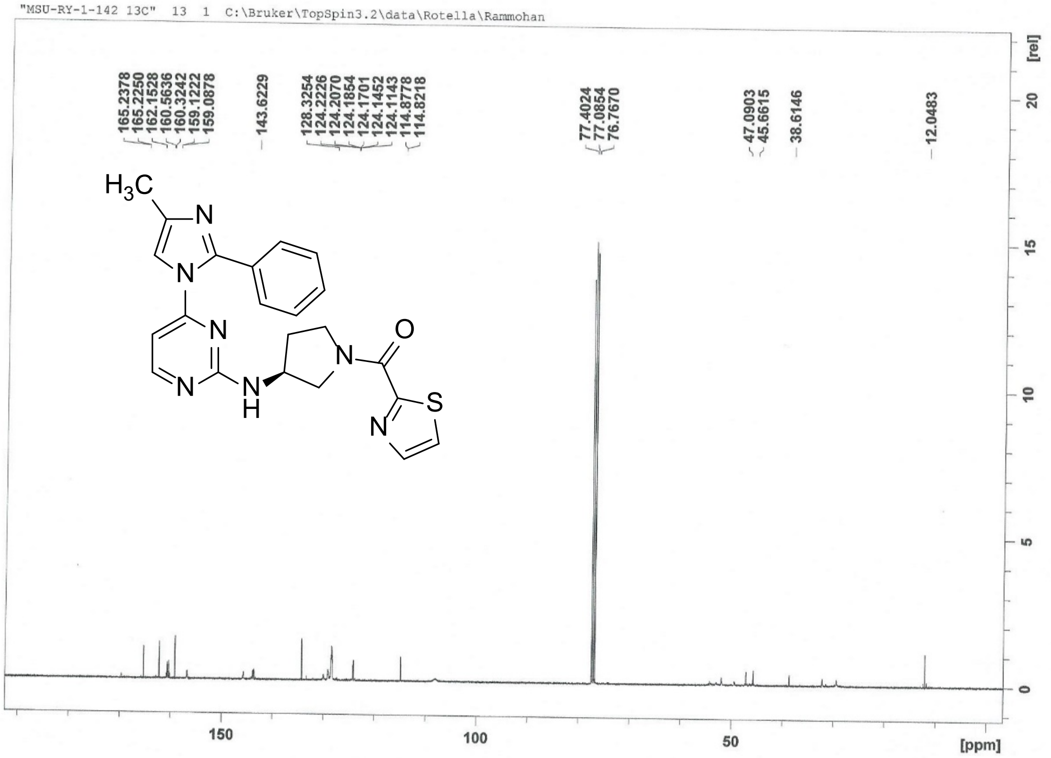


### ==== Shimadzu LabSolutions Data Report ====

#### MSU-RY-1-142

Line#:1 R.Time:2.723(Scan#:----)
MassPeaks:21
RawMode:Averaged 2.722-2.725(1614-1616) BasePeak:432(10016)
BG Mode:Calc Segment 1 - Event 1

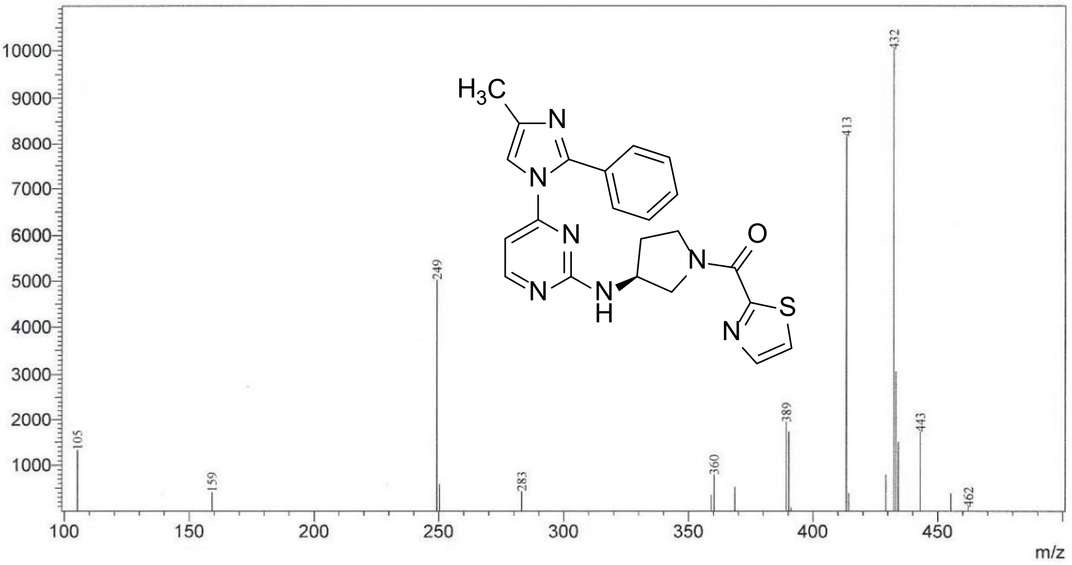

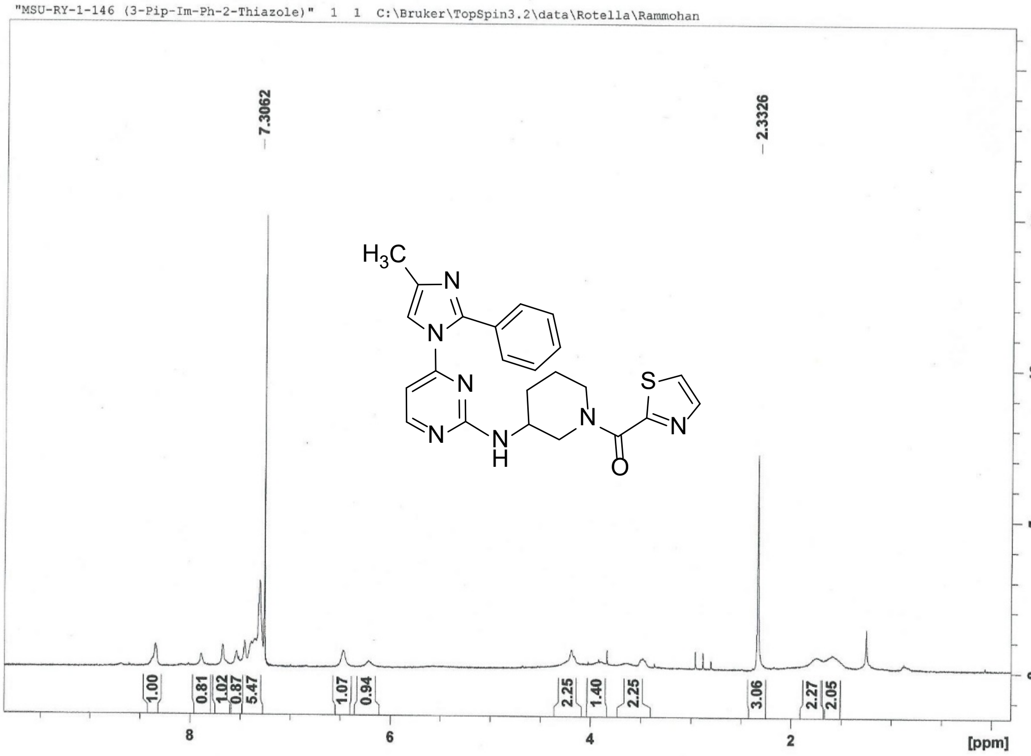
"MSU-RY-1-146 (3-Pip-Im-Ph-2-Thiazole)" 1 1 C:\Bruker\TopSpin3.2\data\Rotella\Rammohan
7.3062
2.3326
1.00
0.81
1.02
0.87
5.47
1.07
0.94
2.25
1.40
2.25
3.06
2.27
2.05
[ppm]
[rel]

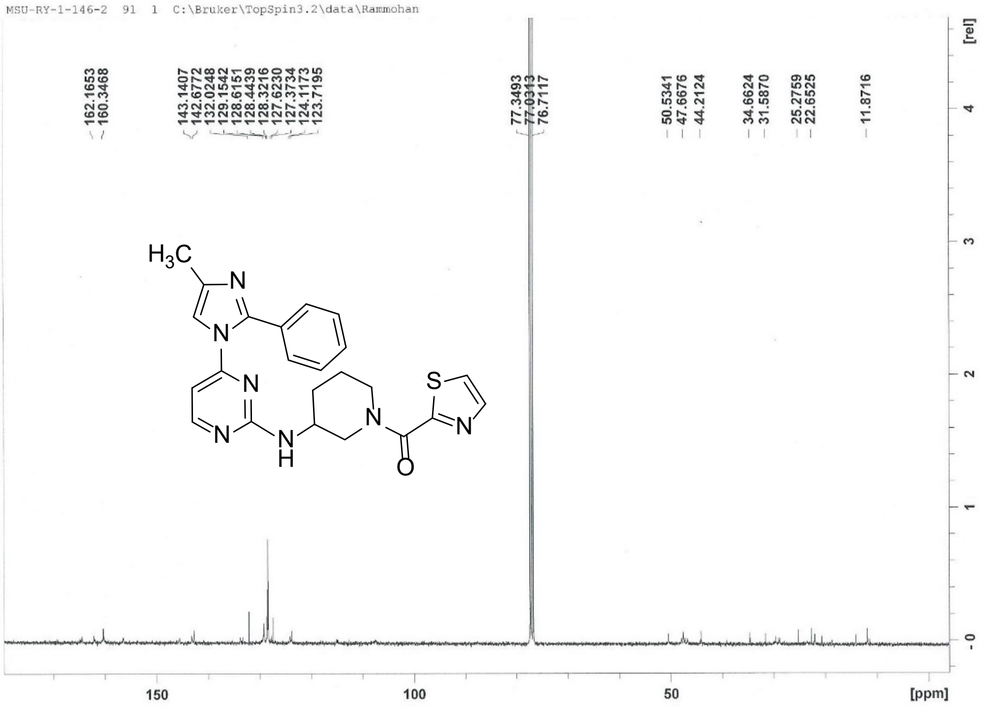


### ==== Shimadzu LabSolutions Data Report ====

**MSU-RY-1-146**

Line#:1 R.Time:2.787(Scan#:1653)
MassPeaks:7
RawMode:Single 2.787(1653) BasePeak:446(6721)
BG Mode:None Segment 1 - Event 1

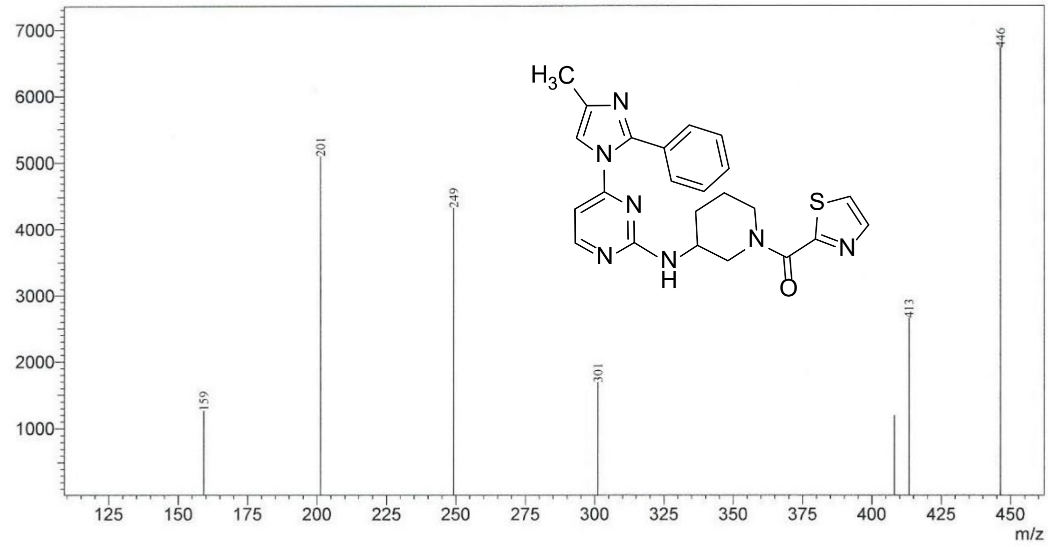

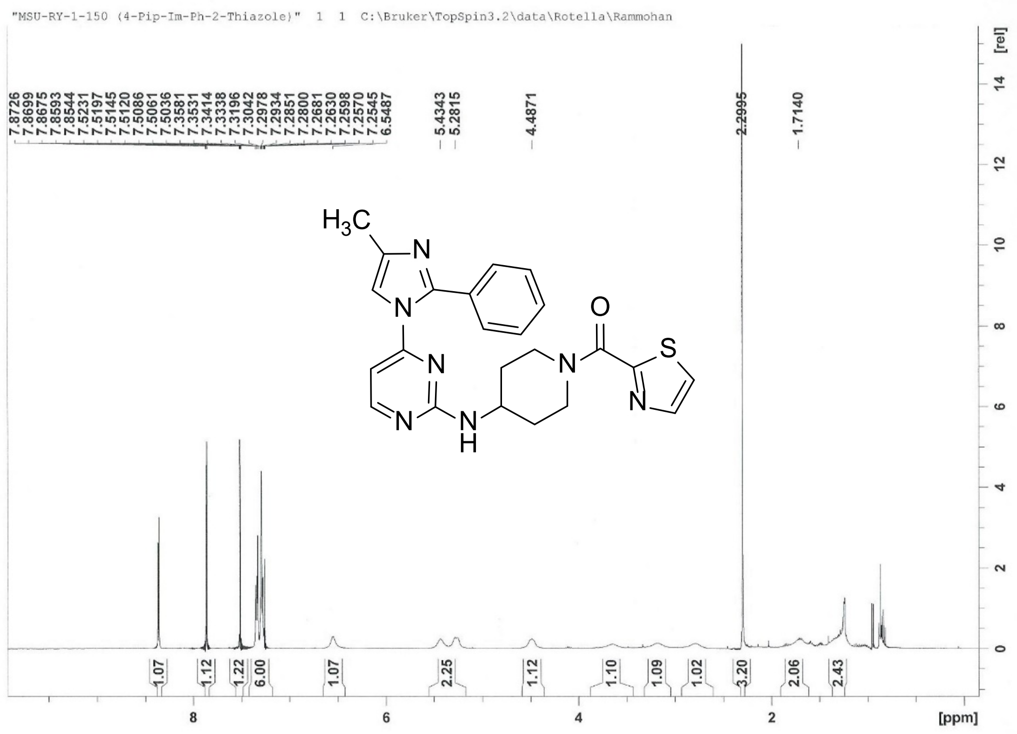
"MSU-RY-1-150 (4-Pip-Im-Ph-2-Thiazole)" 1 1 C:\Bruker\TopSpin3.2\data\Rotella\Rammohan
7.8726
7.8699
7.8675
7.8593
7.8544
7.5231
7.5197
7.5145
7.5120
7.5086
7.5061
7.5036
7.3581
7.3531
7.3414
7.3338
7.3196
7.3042
7.2978
7.2934
7.2851
7.2800
7.2681
7.2630
7.2598
7.2570
7.2545
6.5487
5.4343
5.2815
4.4871
2.2995
1.7140
1.07
1.12
1.22
6.00
1.07
2.25
1.12
1.10
1.09
1.02
3.20
2.06
2.43
[ppm]
[rel]

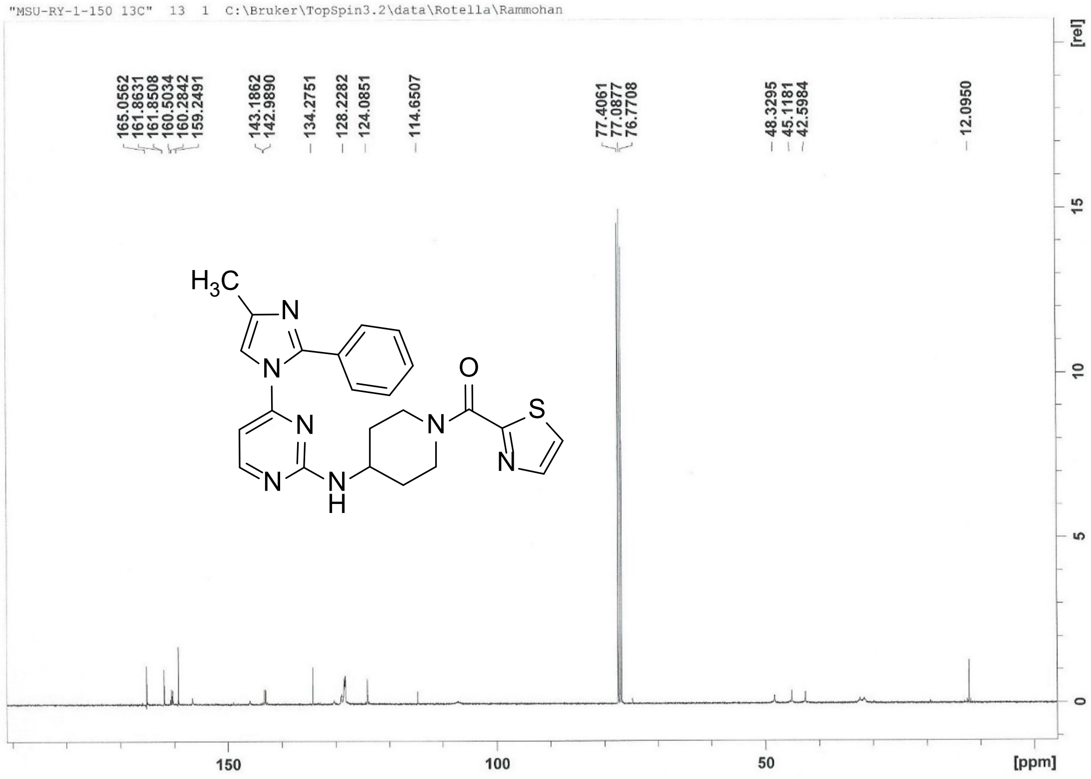
"MSU-RY-1-150 13C" 13 1 C:\Bruker\TopSpin3.2\data\Rotella\Rammohan
165.0562
161.8631
161.8508
160.5034
160.2842
159.2491
143.1862
142.9890
134.2751
128.2282
124.0851
114.6507
77.4061
77.0877
76.7708
48.3295
45.1181
42.5984
12.0950
[rel]
[ppm]

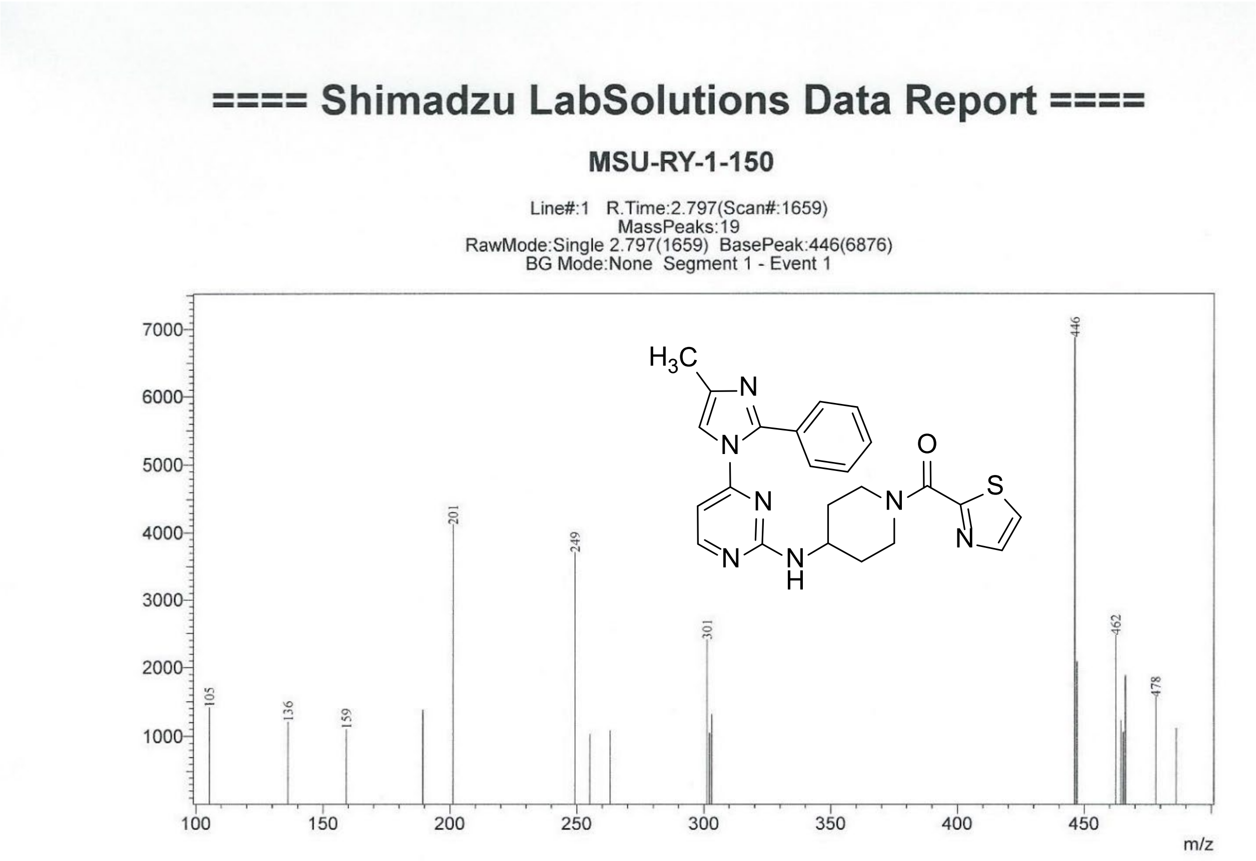


**S1.6.** $^{1}$H, $^{13}$C NMR and Mass spectra of **MSU-RY-1-165**

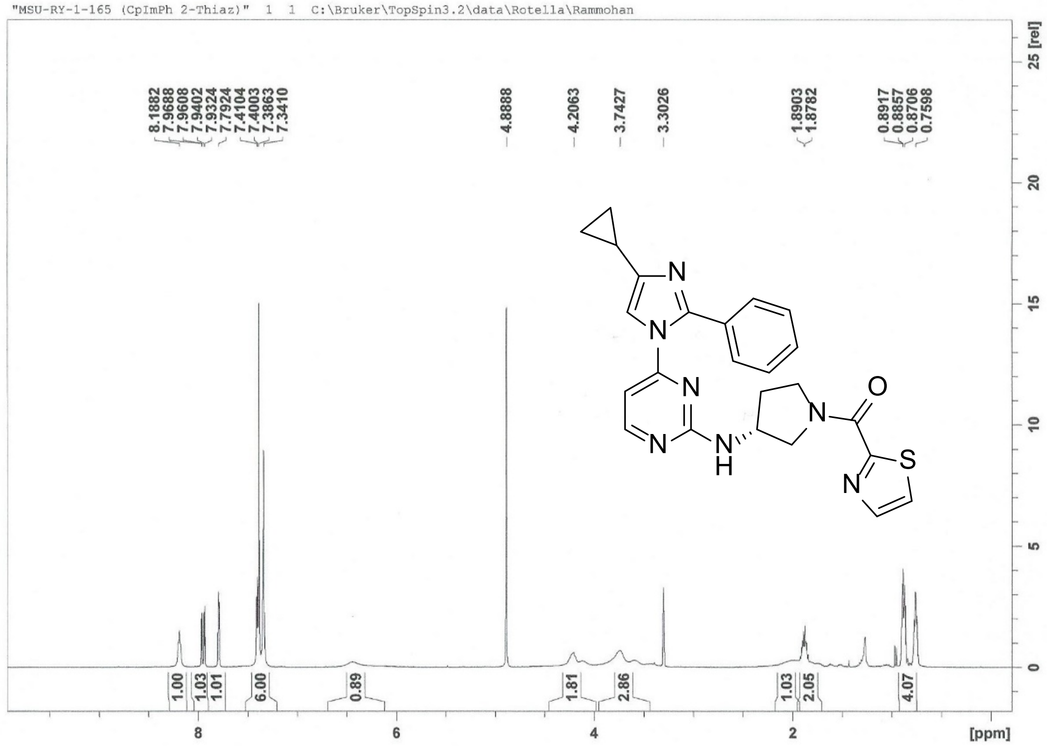
"MSU-RY-1-165 (CpImPh 2-Thiaz)" 1 1 C:\Bruker\TopSpin3.2\data\Rotella\Rammohan
8.1882
7.9688
7.9608
7.9402
7.9324
7.7924
7.4104
7.4003
7.3863
7.3410
4.8888
4.2063
3.7427
3.3026
1.8903
1.8782
0.8917
0.8857
0.8706
0.7598
1.00
1.03
1.01
6.00
0.89
1.81
2.86
1.03
2.05
4.07
[ppm]
25 [rel]
N
N
N
N
H
N
O
N
S
N

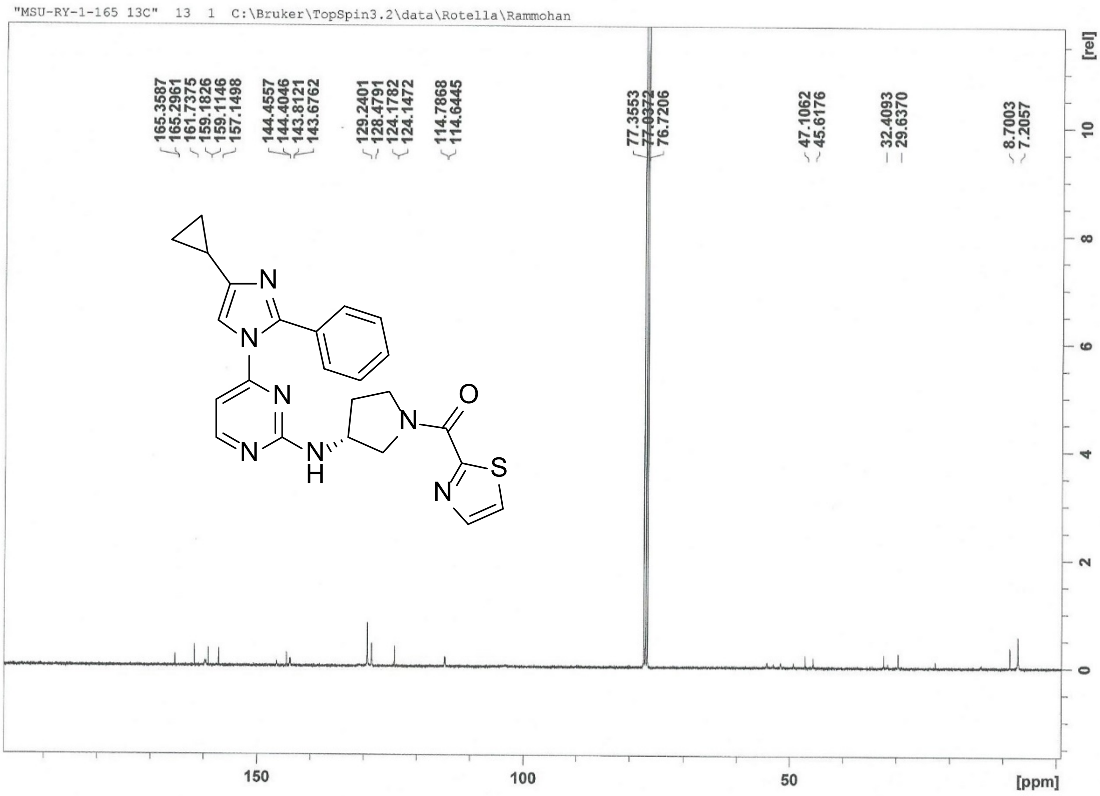


### ==== Shimadzu LabSolutions Data Report ====

#### MSU-RY-1-165

Line#:1 R.Time:2.745(Scan#:----)
MassPeaks:9
RawMode:Averaged 2.743-2.747(1627-1629) BasePeak:458(183509)
BG Mode:Calc Segment 1 - Event 1

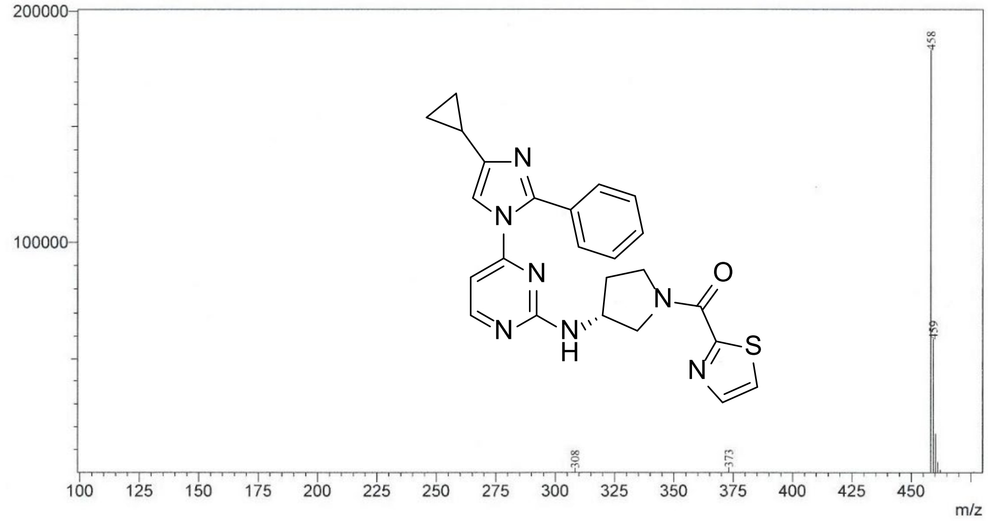

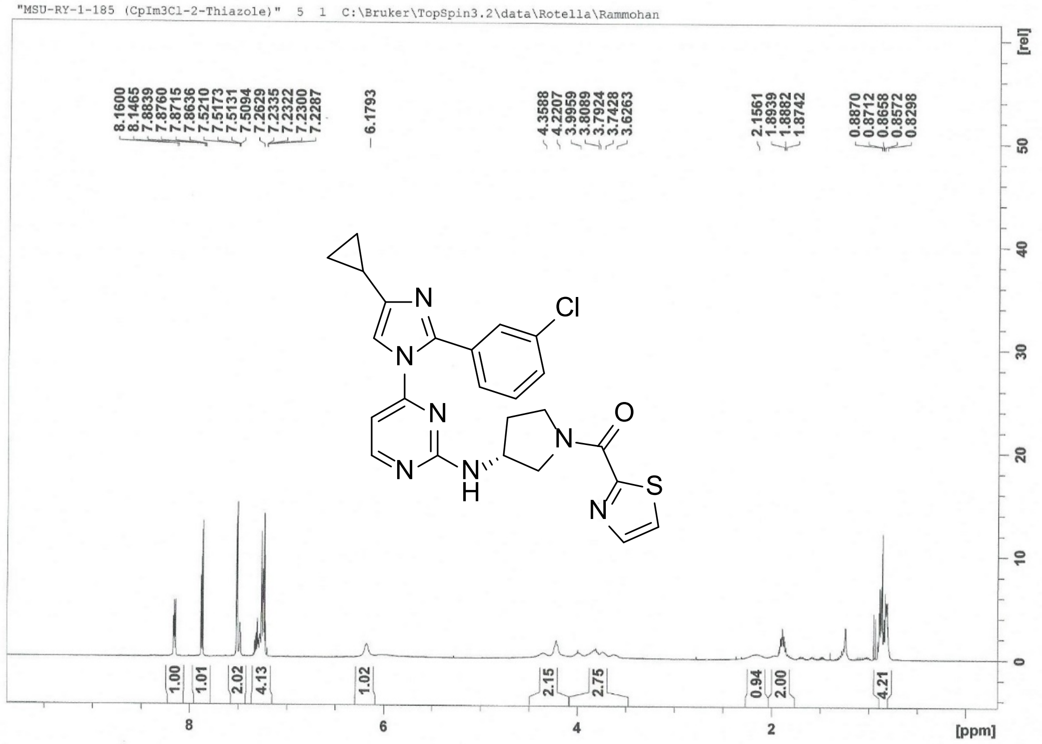
"MSU-RY-1-185 (CpIm3Cl-2-Thiazole)" 5 1 C:\Bruker\TopSpin3.2\data\Rotella\Rammohan
8.1600
8.1465
7.8839
7.8760
7.8715
7.8636
7.5210
7.5173
7.5131
7.5094
7.2629
7.2335
7.2322
7.2300
7.2287
6.1793
4.3588
4.2207
3.9959
3.8089
3.7924
3.7428
3.6263
2.1561
1.8939
1.8882
1.8742
0.8870
0.8712
0.8658
0.8572
0.8298
N
Cl
N
N
O
N
N
N
S
H
N
1.00
1.01
2.02
4.13
1.02
2.15
2.75
0.94
2.00
4.21
[rel]
[ppm]

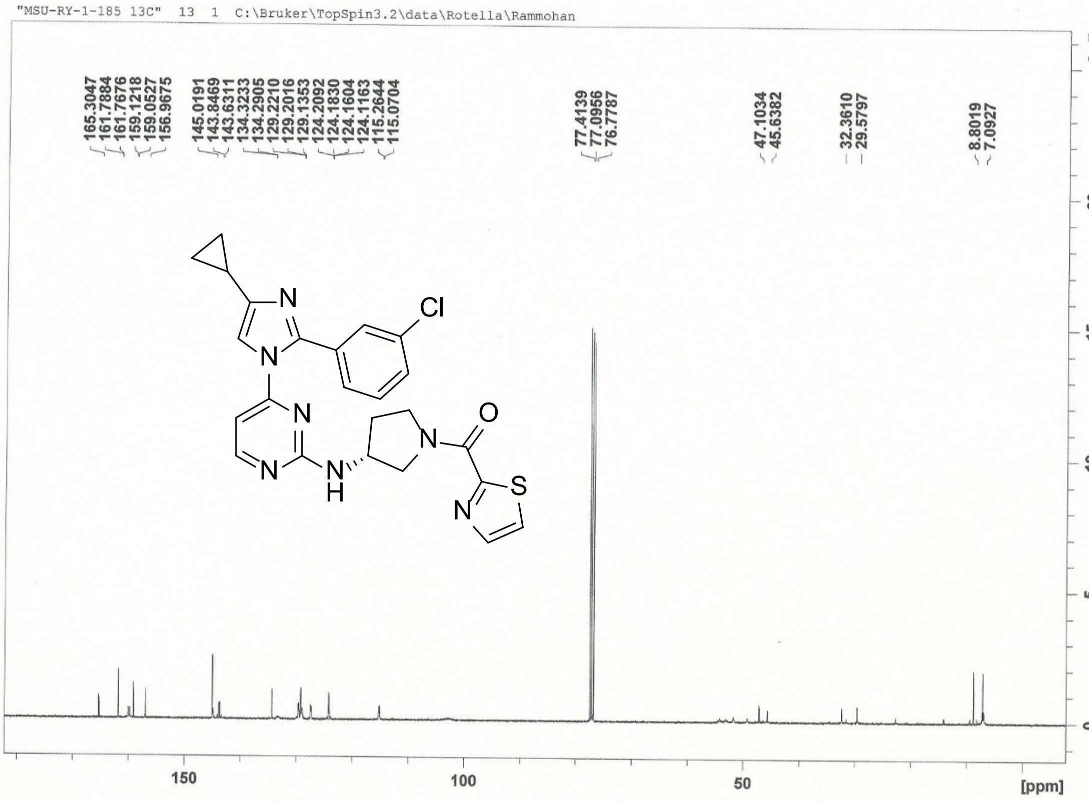
"MSU-RY-1-185 13C" 13 1 C:\Bruker\TopSpin3.2\data\Rotella\Rammohan
165.3047
161.7884
161.7676
159.1218
159.0527
156.9675
145.0191
143.8469
143.6311
134.3233
134.2905
129.2210
129.2016
129.1353
124.2092
124.1830
124.1604
124.1163
115.2644
115.0704
77.4139
77.0956
76.7787
47.1034
45.6382
32.3610
29.5797
8.8019
7.0927
N
Cl
N
N
O
N
N
N
S
H
N
[rel]
[ppm]

### ==== Shimadzu LabSolutions Data Report ====

#### MSU-RY-1-185

Line#:1 R.Time:3.330(Scan#:1979)
MassPeaks:3
RawMode:Single 3.330(1979) BasePeak:492(3151)
BG Mode:None Segment 1 - Event 1

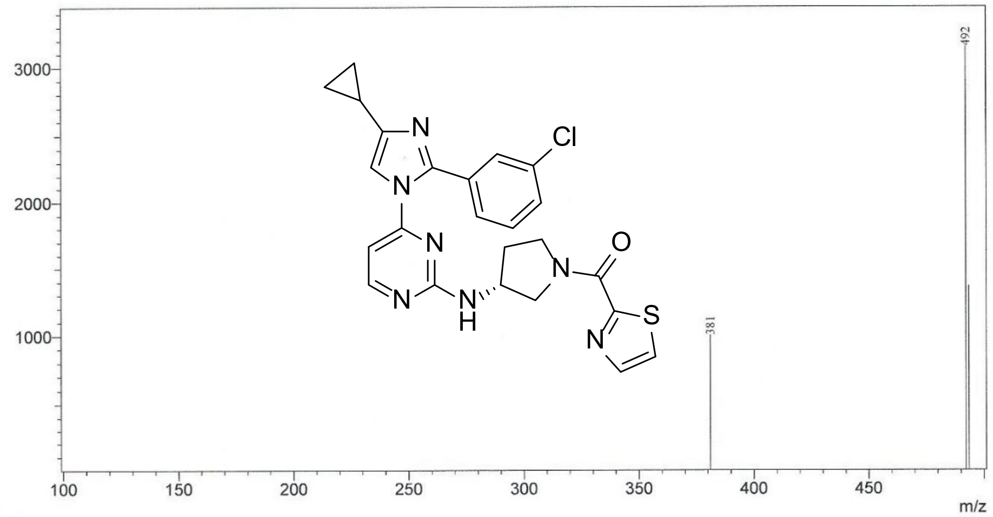


"MSU-RY-2-21-1 (4-Me-2-PhOMe 2-Thiazole)" 5 1 C:\Bruker\TopSpin3.2\data\Rotella\Rammohan

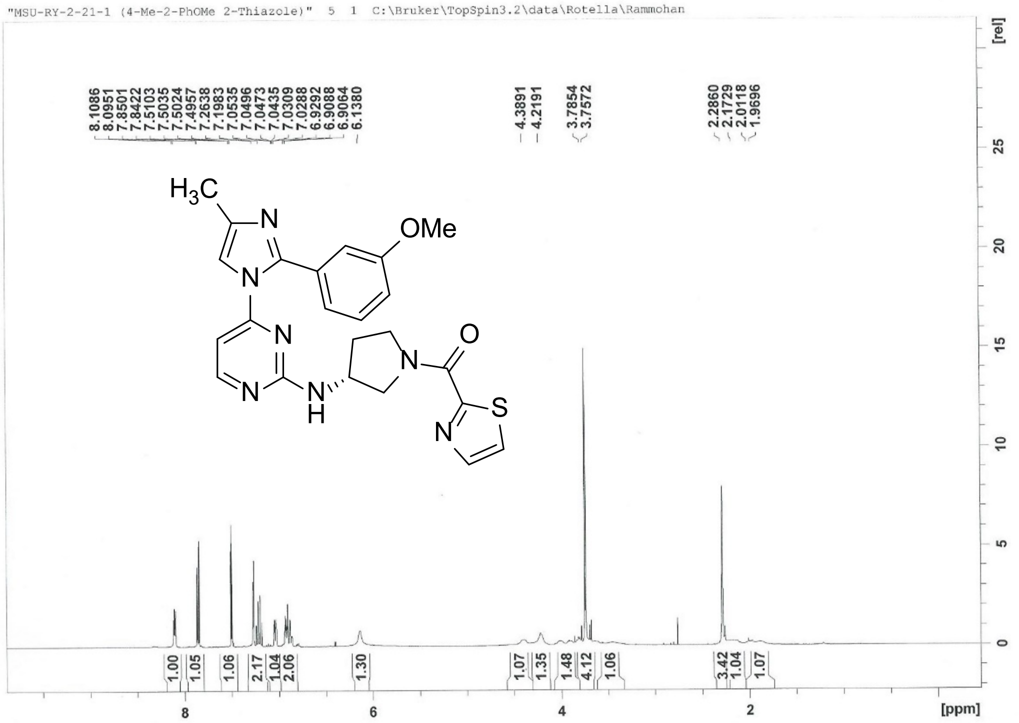

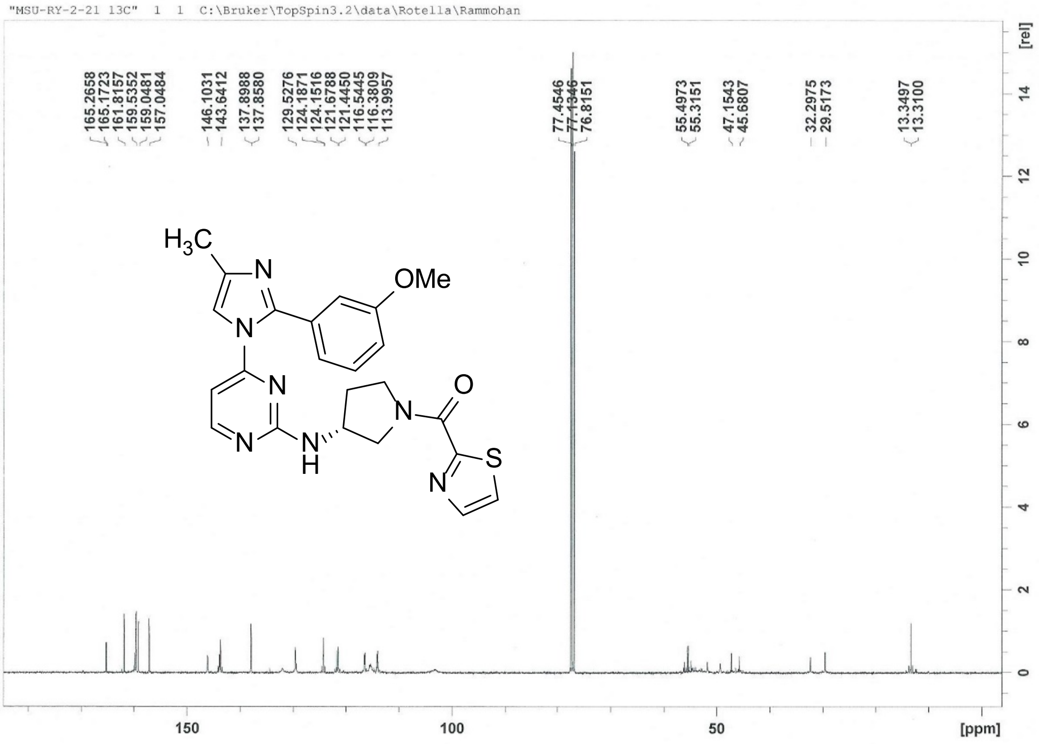


### ==== Shimadzu LabSolutions Data Report ====

#### MSU-RY-2-21

Line#:1 R.Time:2.525(Scan#:----)
MassPeaks:19
RawMode:Averaged 2.523-2.527(1495-1497) BasePeak:462(163631)
BG Mode:Calc Segment 1 - Event 1

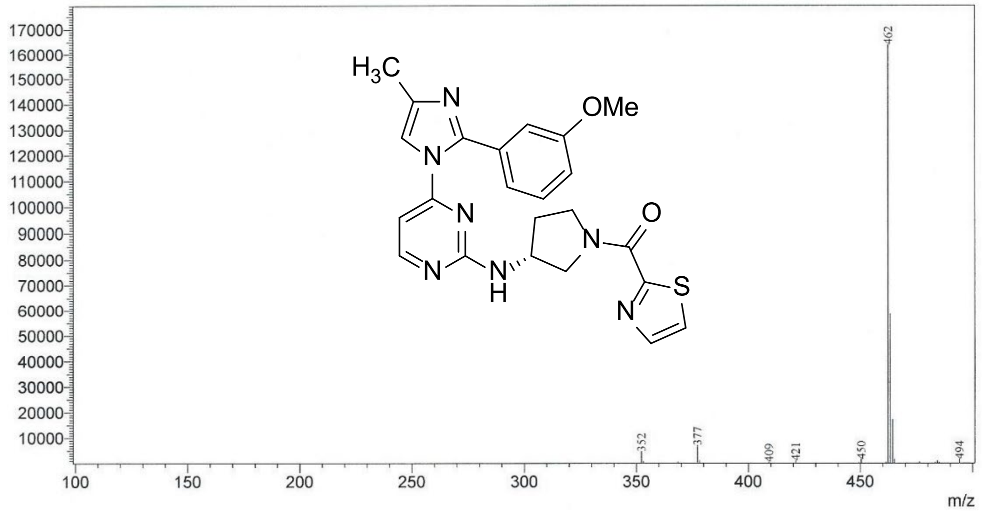

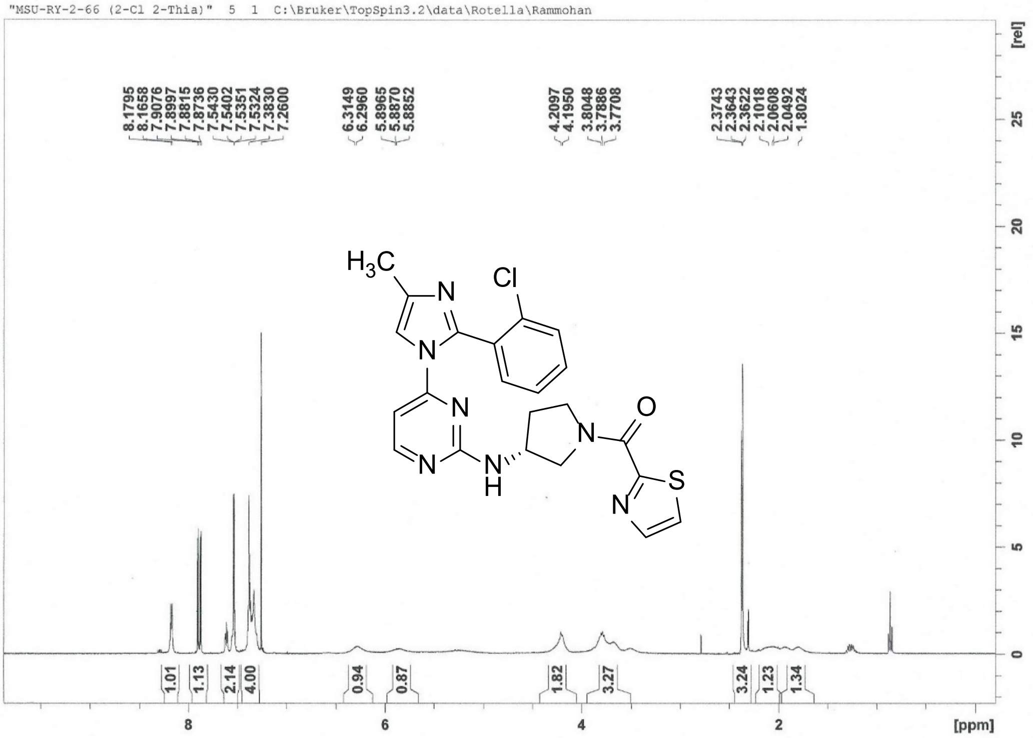

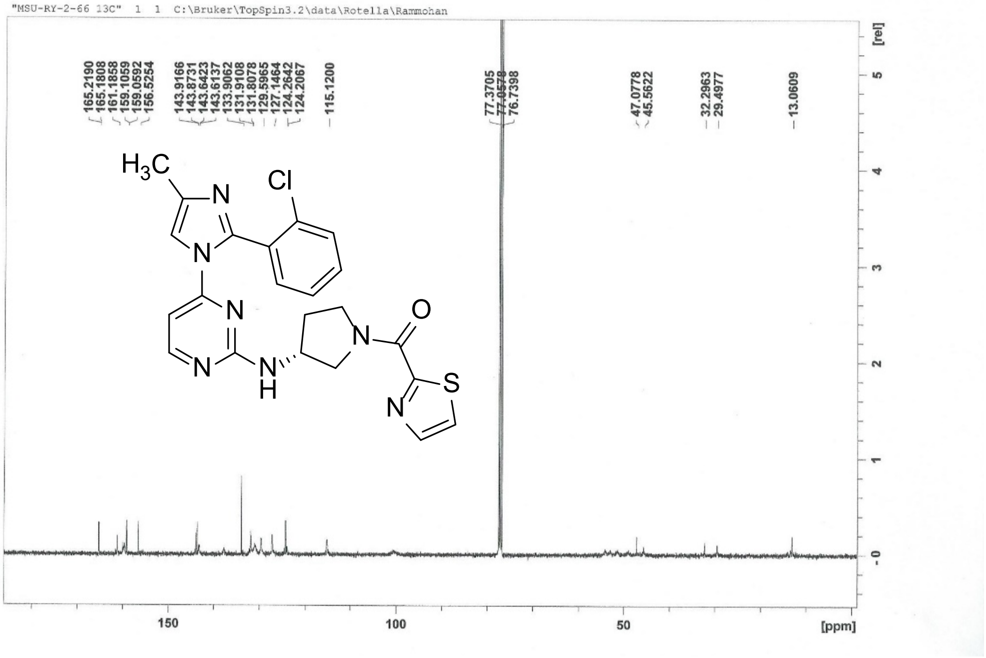

==== Shimadzu LabSolutions Data Report ====

**MSU-RY-2-66**

Line#:1 R.Time:3.000(Scan#:1781)
MassPeaks:3
RawMode:Single 3.000(1781) BasePeak:466(3041)
BG Mode:None Segment 1 - Event 1

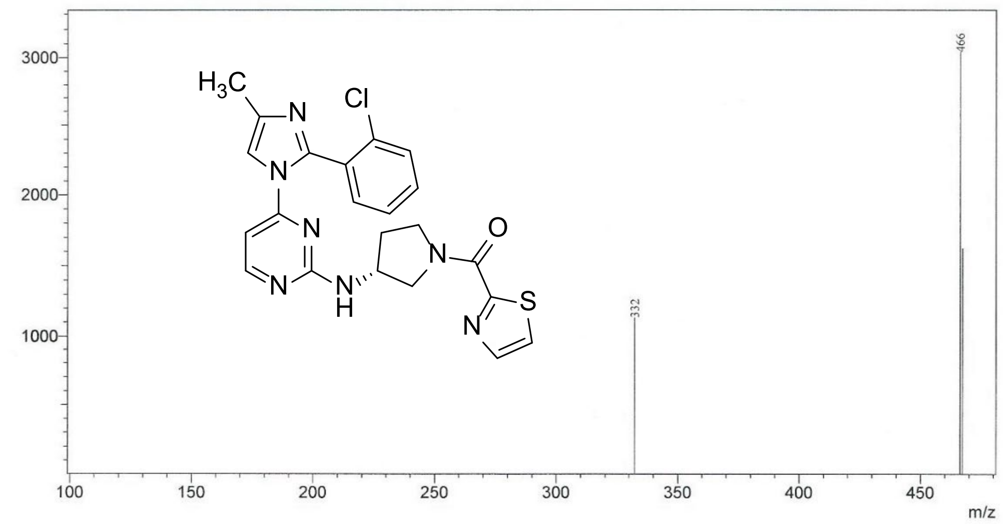


"MSU-RY-2-68 (4-Cl 2-Thia)" 5 1 C:\Bruker\TopSpin3.2\data\Rotella\Lozano

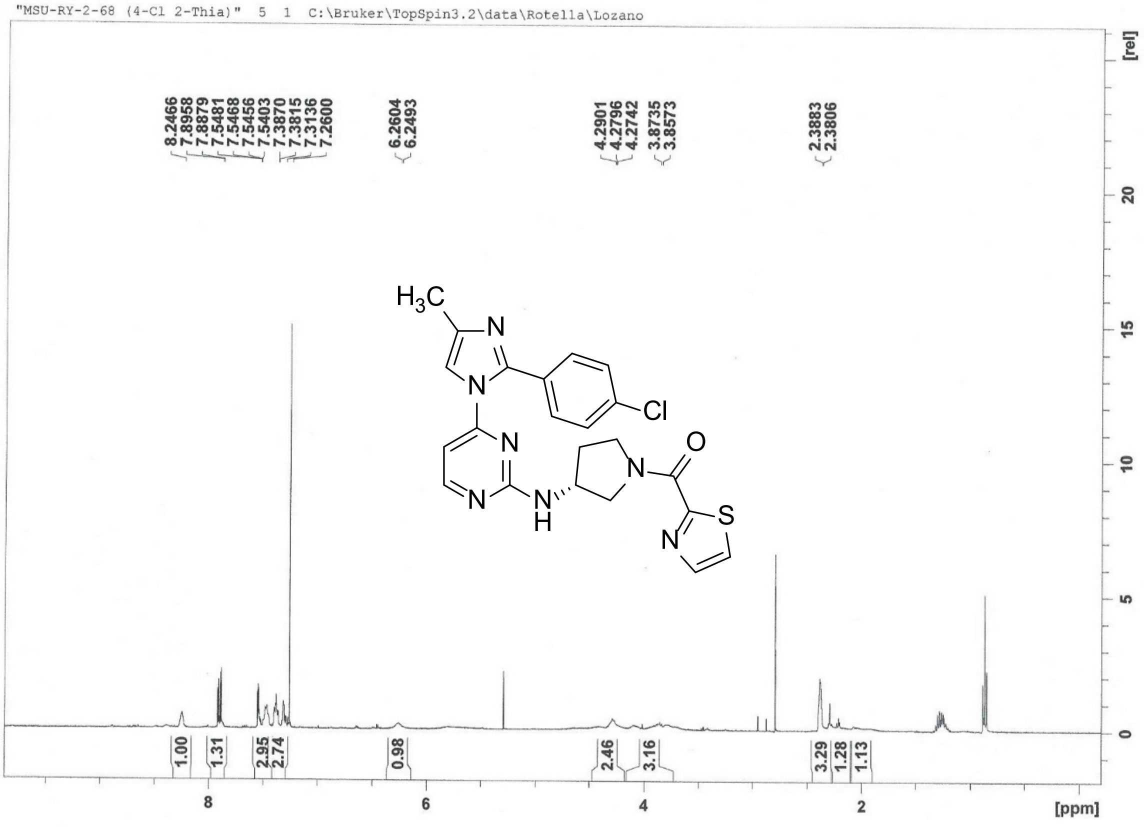

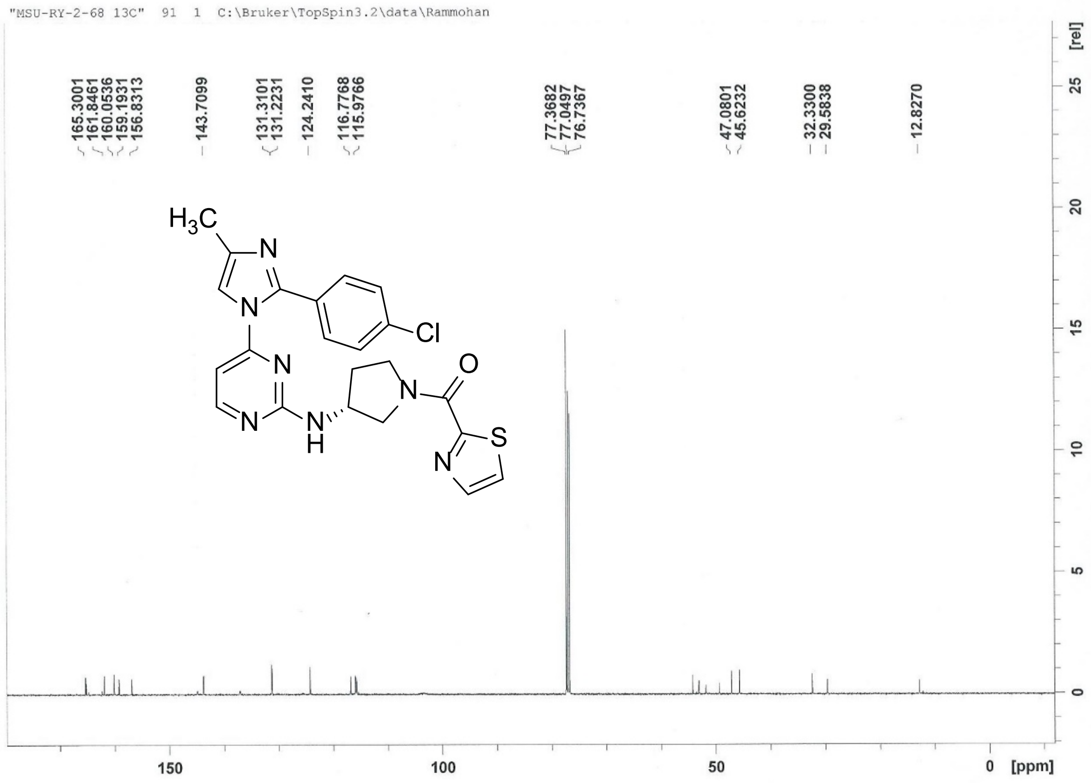
"MSU-RY-2-68 13C" 91 1 C:\Bruker\TopSpin3.2\data\Rammohan
165.3001
161.8461
160.0536
159.1931
156.8313
143.7099
131.3101
131.2231
124.2410
116.7768
115.9766
77.3682
77.0497
76.7367
47.0801
45.6232
32.3300
29.5838
12.8270
H3C
N
N
Cl
N
N
N
O
N
H
S
N
[rel]
25
20
15
10
5
0
150
100
50
0
[ppm]

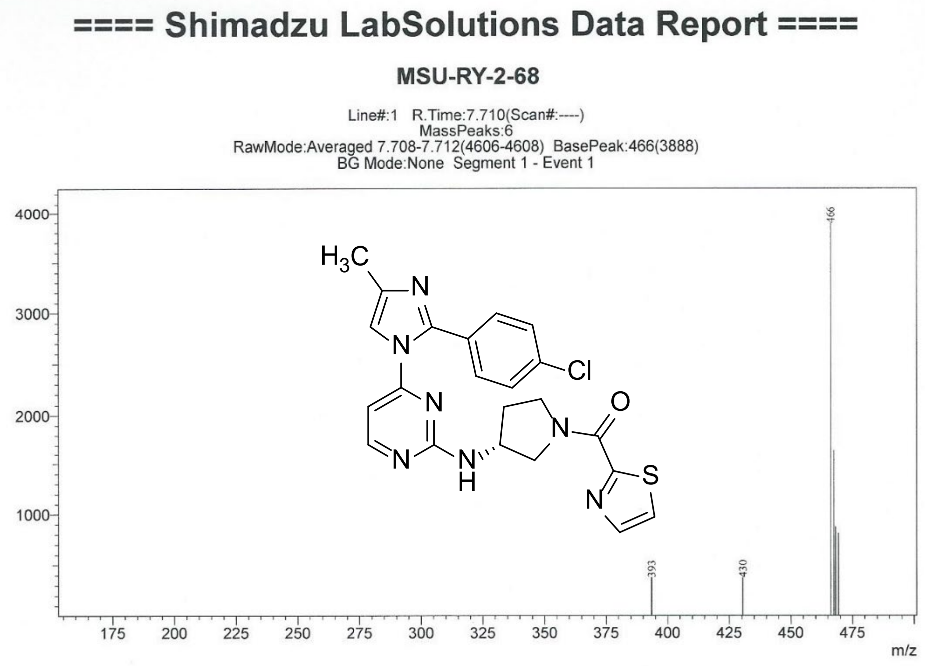
==== Shimadzu LabSolutions Data Report ====
MSU-RY-2-68
Line#:1 R.Time:7.710(Scan#:----)
MassPeaks:6
RawMode:Averaged 7.708-7.712(4606-4608) BasePeak:466(3888)
BG Mode:None Segment 1 - Event 1
4000
3000
2000
1000
466
393
430
H3C
N
N
Cl
N
N
N
O
N
H
S
N
175
200
225
250
275
300
325
350
375
400
425
450
475
m/z

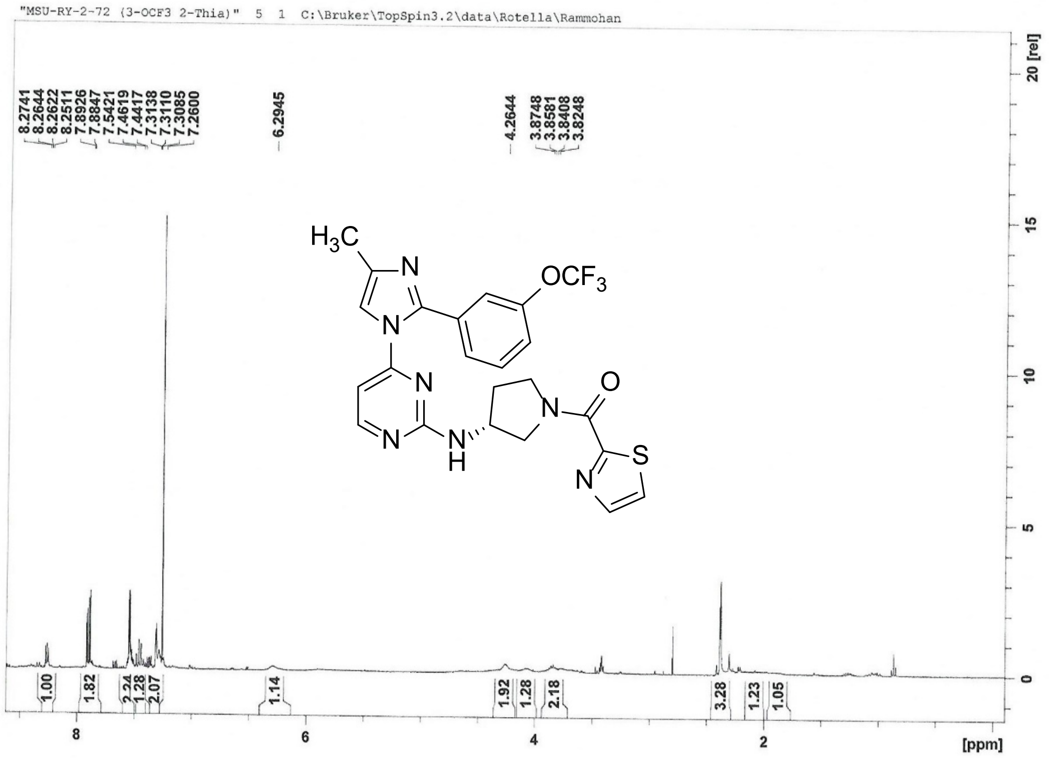
"MSU-RY-2-72 (3-OCF3 2-Thia)" 5 1 C:\Bruker\TopSpin3.2\data\Rotella\Rammohan
8.2741
8.2644
8.2622
8.2511
7.8926
7.8847
7.5421
7.4619
7.4417
7.3138
7.3110
7.3085
7.2600
6.2945
4.2644
3.8748
3.8581
3.8408
3.8248
$H_3C$
N
$OCF_3$
N
N
O
N
N
N
S
H
N
1.00
1.82
2.24
1.28
2.07
1.14
1.92
1.28
2.18
3.28
1.23
1.05
[ppm]
[rel]

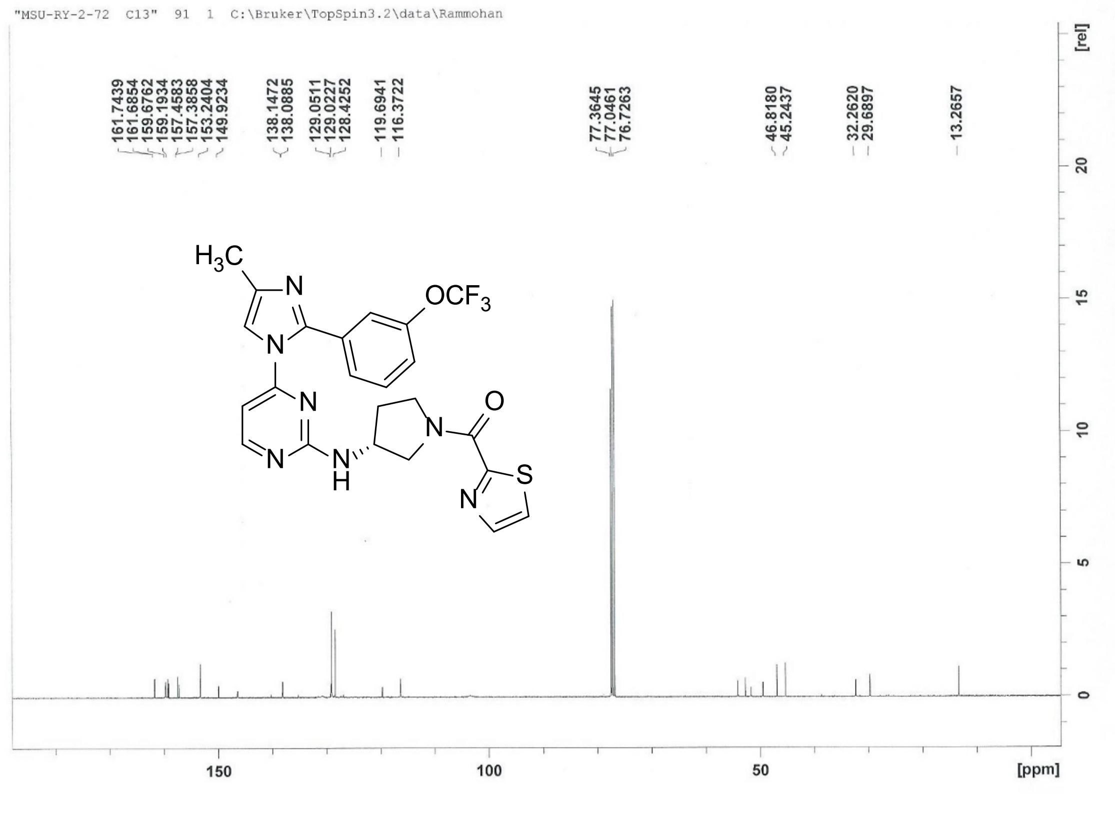
"MSU-RY-2-72 C13" 91 1 C:\Bruker\TopSpin3.2\data\Rammohan
161.7439
161.6854
159.6762
159.1934
157.4583
157.3858
153.2404
149.9234
138.1472
138.0885
129.0511
129.0227
128.4252
119.6941
116.3722
77.3645
77.0461
76.7263
46.8180
45.2437
32.2620
29.6897
13.2657
$H_3C$
N
$OCF_3$
N
N
O
N
N
N
S
H
N
[ppm]
[rel]

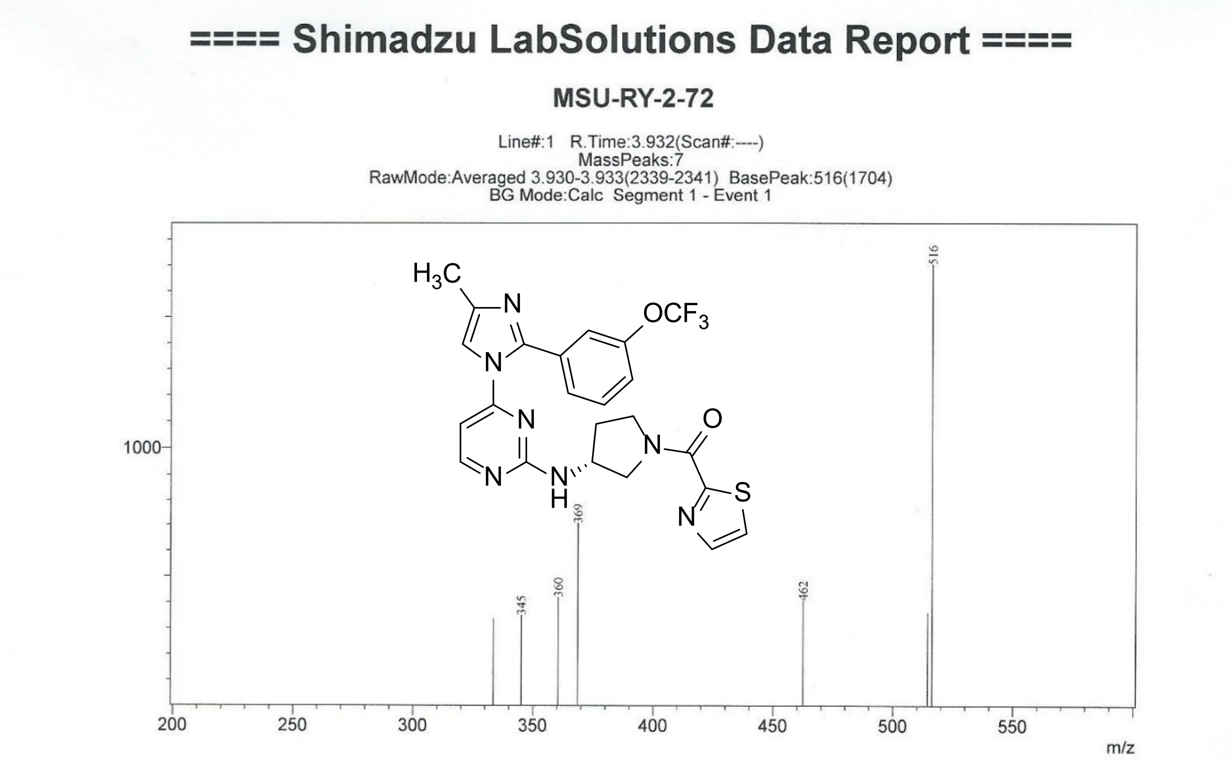


#### PfPKG Enzymatic Assays

##### $IC_{50}$ determination

$IC_{50}$ values were determined using the commercial immobilized metal ion affinity-based fluorescence polarization (IMAP) assay (Molecular Devices # R8127). In a 20 µL reaction, the following amounts of enzymes were used in each assay well: 17 ng of wild type PfPKG, 7 ng of T618Q PfPKG, and 10 ng of hPKG. Enzymes were preincubated at room temperature for 15 minutes with inhibitor concentrations ranging from 4 nM to 10 µM in assay buffer RB-T (10 mM Tris-HCl, pH 7.2, 10 mM $MgCl_2$, 0.05% $NaN_3$, 0.01% Tween®20). Next, 120 nM fluorescent peptide substrate (FAM-PKAtide or FAM-IP3R-derived peptide), 10 µM ATP, 1 µM cGMP, and 1.0 mM DTT were added to each well to begin the reaction. After a 60-minute incubation at room temperature, the reaction was halted and developed with 60 µL of the progressive binding reagent (PBR). For PfPKG and the mutant enzyme, the PBR was diluted 400-fold in 100% 1X IMAP Progressive Binding Buffer A and incubated for 30 minutes. For the human PKG, the PBR was diluted 400-fold in 75% 1X IMAP Progressive Binding Buffer A and 25% 1X IMAP Progressive Binding Buffer B, then incubated for 60 minutes. Fluorescent polarization was read in parallel and perpendicular to the excitation plane (ex. 485 nm/ em. 528 nm) using a Synergy 2 Microplate reader (BioTek, Winooski, VT). Two replicate wells were used for each sample and averaged before further analysis. The data were analyzed using a four-parameter logistic curve using Microsoft Excel Solver to make dose response curves in Microsoft Excel. To compare potency and assay quality, **5** was used as a positive control in each experiment.

#### In vitro ADME

Metabolic stability: Incubate samples at 37°C in the presence of human or mouse liver microsomes and NADPH according to standard methods (1). Aliquots are removed at 5 time points, quenched and analyzed (LCMS/MS and MS/MS as needed) for remaining test compound, along with a positive control. Microsomal protein content is adjusted to give consistent results. Data are reported as half-life and clearance. Assay acceptance criteria is 20% for all standards and 25% for the LLOQ.

Aqueous solubility: Kinetic aqueous solubility is measured by adding ~2 mg samples to pH 7.4 buffered aqueous solutions at 25°C. The mixture is agitated at 25°C for 1 hour, filtered and evaluated by UV and/or LCMSMS analysis. Data are reported as mg/mL and μM concentration.

CYP inhibition: Compounds are assessed for their ability to inhibit the three major human cytochrome P450 enzymes, 3A4, 2D6 and 2C9. Expressed enzymes are used to minimize non-specific binding and membrane partitioning issues (2). The 3A4 assay uses testosterone as a substrate and is analyzed by LC/MS/MS on a Waters TQ instrument using positive or negative electrospray ionization. The 2D6 and 2C9 assays use fluorescent substrates and are analyzed on an Envision Plate Reader.

hERG inhibition: HEK293 cells stably transfected with the hERG ion channel are grown to 80% confluency and then seeded into poly-lysine coated plates (25,000 cells/well). Cells are loaded with Thallos dye, treated with test compounds at 1 and 10 μM (final DMSO concentration = 0.1%) and then thalium flux is measured on a plate reader (excitation @ 480 nM; emission @ 530 nM) according to published procedures (3,4). Data are reported as percent inhibition at both concentrations.

#### Cellular parasite infectivity assays:

##### *In vitro P. cynomolgi* assay

The *P. cynomolgi* liver stage experiment was performed as previously described[5]. Briefly, cryopreserved primary non-human primate hepatocytes (lot KXA) and hepatocyte culture medium (HCM) (InVitroGroTM CP Medium) were obtained from BioIVT, Inc., (Baltimore, MD, USA) and thawed following manufacturer recommendations. The hepatocytes were plated into pre-collagen coated 384-well plates (Cat No. 781956, Greiner, Monroe, NC, U.S.A.) and used for experiments within 2 – 4 days after plating. Infectious sporozoites were obtained from *An. dirus mosquitoes* infected with *P. cynomolgi* B strain and diluted accordingly (600 sporozoites/μL) in HCM. Compounds were dissolved in 100% DMSO and used at a final starting concentration of 20 μM in a 12-point, 3-fold serial dilution. The compounds were dosed in two treatment modes, prophylactic and radical cure. In prophylactic mode, drug was present for 4 days starting at point of sporozoite addition. Alternatively, in radical cure mode, drug was present for 4 days starting on day 4 post sporozoite inoculation.

Imaging and data analysis of the drug plates were completed using the Operetta CLS Imaging System and Harmony software 4.16 (Perkin Elmer, Waltham, MA, USA). Images were acquired using TRITC, DAPI, and bright field channels using a 10x objective. Following methodology previously described[6], hepatocyte health (toxicity) and parasite populations (schizonts and hypnozoites) were identified and quantified by the specific object properties including area, mean intensity, maximum intensity, and cell roundness. Percent inhibition was calculated using the following equation, % inhibition = 100 x [(X – AVG $N_{ctrl}$) / (AVG $P_{ctrl}$ – AVG $N_{ctrl}$)] where X is the replicate average at the specific compound concentration, AVG $P_{ctrl}$ is the replicate average of the assay positive control (tafenoquine), and AVG Nctrl is the replicate average of the assay negative control (DMSO). The percent inhibition values were used to calculate the half maximal inhibitory concentration ($IC_{50}$) and dose-response modeling was performed in GraphPad Prism version 8.1.0 (GraphPad, La Jolla, CA, USA) using the log(inhibitor) vs. response -- Variable slope (four parameters) model. The reported $IC_{50}$ values are expressed as mean percent inhibition ± standard deviation (S.D.) from experimental replicates (n=2) with biological replicates (n=2) for both drug treatment modes.

***P. berghei*** **sporozoite infectivity assay:**

HepG2 cells were plated in 8-well LabTek chamber slides were infected with approximately 2 x $10^4$ luciferase-expressing *P. berghei* sporozoites in the presence of compounds or vehicle. Compound-containing media was replaced with compound-free medium at 16 post-infection. At 48 hours post-infection, medium was replaced with D-luciferin diluted in PBS. Luciferase activity was measured using an IVIS® Spectrum in vivo imaging system.

5. Vanachayangkul, P.; Im-Erbsin, R.; Tungtaeng, A.; Kodchakorn, C.; Roth, A.; Adams, J.; Chaisatit, C.; Saingam, P.; Sciotti, R. J.; Reichard, G. A.; Nolan, C. K.; Pybus, B. S.; Black, C. C.; Lugo-Roman, L. A.; Wegner, M. D.; Smith, P. L.; Wojnarski, M.; Vesely, B. A.; Kobylinski, K. C., Safety, Pharmacokinetics, and Activity

of High-Dose Ivermectin and Chloroquine against the Liver Stage of Plasmodium cynomolgi Infection in Rhesus Macaques. *Antimicrob Agents Chemother* **2020,** *64* (9).

6. Roth, A.; Maher, S. P.; Conway, A. J.; Ubalee, R.; Chaumeau, V.; Andolina, C.; Kaba, S. A.; Vantaux, A.; Bakowski, M. A.; Thomson-Luque, R.; Adapa, S. R.; Singh, N.; Barnes, S. J.; Cooper, C. A.; Rouillier, M.; McNamara, C. W.; Mikolajczak, S. A.; Sather, N.; Witkowski, B.; Campo, B.; Kappe, S. H. I.; Lanar, D. E.; Nosten, F.; Davidson, S.; Jiang, R. H. Y.; Kyle, D. E.; Adams, J. H., A comprehensive model for assessment of liver stage therapies targeting Plasmodium vivax and Plasmodium falciparum. *Nat Commun* **2018,** *9* (1), 1837.
